# The molecular architecture of the Arabidopsis callose synthase complex

**DOI:** 10.64898/2026.05.05.722910

**Authors:** David Ušák, Petra Cifrová, Michal Daněk, Tereza Korec Podmanická, Daniela Kocourková, Dominique Eeckhout, Jelle Van Leene, Adam Zeiner, Michaela Neubergerová, Julia M. Kraus, Roman Hudeček, Judith Garcia Gonzalez, Anastasiia Zhivaeva, Anzhela Antonova, Tetiana Kalachova, Geert De Jaeger, Michael Wrzaczek, Daniel Van Damme, Roman Pleskot

**Affiliations:** Institute of Experimental Botany, Czech Academy of Sciences, 16502 Prague, Czech Republic; Department of Experimental Plant Biology, Faculty of Science, 12000 Charles University, Prague, Czech Republic; Department of Plant Biotechnology and Bioinformatics, Ghent University, Technologiepark 71, 9052 Ghent, Belgium; VIB Center for Plant Systems Biology, Technologiepark 71, 9052 Ghent, Belgium; Institute of Plant Molecular Biology, Biology Centre, Czech Academy of Sciences, 370 05 České Budějovice, Czech Republic; Faculty of Science, University of South Bohemia, 370 05 České Budějovice, Czech Republic

**Keywords:** callose, callose synthase, β-1, 3-glucan, protein structure, evolution, plasmodesmata, molecular dynamics

## Abstract

Callose synthase is responsible for the targeted deposition of the β-1,3-glucan polymer, callose which underlines essential plant developmental processes, including cell division, pathogen defense or cell-cell communication. The architecture of the callose synthase complex (CALSC) as well as the molecular mechanisms of callose synthesis remain unknown. Here we report an integrative characterisation of the Arabidopsis CALS complex, with the most enriched subunits, CALS1, CALS2 and CALS3, forming its core. Structurally, CALSC assembles into a trimer, requiring the plant-specific Bag domain to mediate inter-subunit associations. The biological importance of CALSC assembly is highlighted by the simultaneous loss of CALS1 and CALS3, which abolishes plasmodesmal callose deposition and affects symplastic transport. Site-directed mutagenesis and molecular dynamics simulations depict the topology of the CALS1 active site in detail, including the components of the enzymatic reaction. We pinpoint the translocating tunnel through which the nascent glucan is delivered and mechanistically confirm the role of transmembrane helix 8 in regulating glucan export. Our work provides unprecedented insight into the molecular architecture of the CALSC and the distinct changes from maturation to activity at the plasma membrane, while showcasing the mechanisms involved in callose synthesis at the molecular level.

## INTRODUCTION

The β-1,3-D glucan callose is the most prominent locally-deposited polysaccharide of the plant cell wall (CW). Callose is composed of β-1,3-D linked glucose molecules forming a linear chain, with an incidental occurrence of β-1,6 branches. During plant ontogeny, callose synthesis is essential for a plethora of processes, including cytokinesis^1,2^, regulation of cell-cell communication by plasmodesmata (PD)^3,4^, phloem development^5^, pollen maturation and growth^6,7^ or stress responses^8,9^. The production of callose is controlled by members of the *CALLOSE SYNTHASE* (*CALS*) gene family, encoding large 200 kDa integral membrane proteins. Three functional domains are present in CALS - Vta1, FKS1 and a glycosyl transferase (GT) domain. Located in tandem at the N-terminus, the former two have an unknown function. The GT domain, positioned within a large central cytoplasmic loop, likely confers the enzymatic activity of CALS^10–12^.In the budding yeast CALS homolog ScFKS1, specific residues within the GT domain correspond to those that form the active site of bacterial cellulose synthase (CESA)^13,13–16^.

CALS is believed to work as part of a larger assembly, together called the CALS complex (CALSC)^2,10,17–20^, with a size roughly 30 nm across^21^. Recently, the structure of ScFKS1 has been resolved by cryo-electron microscopy (cryo-EM) - its size of 10 nm implies that other proteins will likely be a part of CALSC^13–15^. Numerous studies have attempted to elucidate the interactory network of plant CALSC. It has been proposed that CALS associates with the cytoskeleton, possibly affecting its proper targeting and plasma membrane (PM) activity^20,22,23^. CALS activity requires magnesium^24,25^ and calcium^21,26^, the role of the latter is further substantiated by the ability of annexins to inhibit it^27^. The glucan chain proceeds from the UDP-glucose (UPG) precursors, supplied to the CALSC either by UPG transferase or Sucrose synthase, both known CALS partners^2,3,18^. The UPG transferase (UGT)1-dependent activation possibly relies on its association with ROP1^28^. Indeed, its yeast homolog Rho1p binds the active site of FKS1 and stimulates β-1,3-glucan synthesis^13,29^. Despite these results, the CALS regulatory mechanisms are so far only poorly investigated in plants and the portfolio of CALS interactors to this date remains scarce.

In this report, we used affinity purification and proximity biotinylation to describe the interactome of *Arabidopsis* CALS1. Combined with cross-linking mass spectrometry (XL-MS) and protein structure prediction, we propose a structural model of the plant CALSC. Our model is further supported by the genetic analysis of *cals1*/*cals3* mutant plants and analysis of CALS1 localization. We uncover the shared cellular roles of CALS1 and CALS3 as CALSC components required for the callose deposition at PD and describe the developmental defects of plants defective in PD closure. Furthermore, we provide insight into the regulation of CALS activity at the molecular level, as well as the transport of the newly-formed glucan chain into the extracellular space.

## RESULTS AND DISCUSSION

### The CALS1 interactome reveals the complex interplay behind callose synthesis

To uncover the regulatory network of CALS1, we generated an *Arabidopsis* cell culture-based interactome. In order to target both stable and transient interactors, we utilized GS^rhino^ pull down along with TurboID proximity labeling, identifying 1350 and 937 unique proteins, respectively (**Supplementary data**). To reduce the background of the analysis, we performed filtering against in-house AP-MS and TurboID-MS control datasets, followed by spectral count (NSAF)-based fold-change and p-value-based removal of non-specific co-purified proteins. The final dataset of significantly enriched proteins contained 220 proteins found by pull down and 104 by TurboID (**Fig. 1a**). Out of these, 24 proteins were identified by both methods, which included the three most enriched proteins across both datasets (CALS1, CALS2 and CALS3), CALS homolog CALS10, lipid-associated proteins (ACBP3, SMT2, SAC7, MSBP2), protein glycosylation enzymes (ALG9, STT3A), endomembrane transport proteins (CASP, ERDJ2A and SYP81) or CW component-synthesizing enzymes (UXS1, QUA3) as well as the putative PD protein MCTP16 and a small GTPase RABH1b (**Fig. 1a**).

**Fig. 1:**
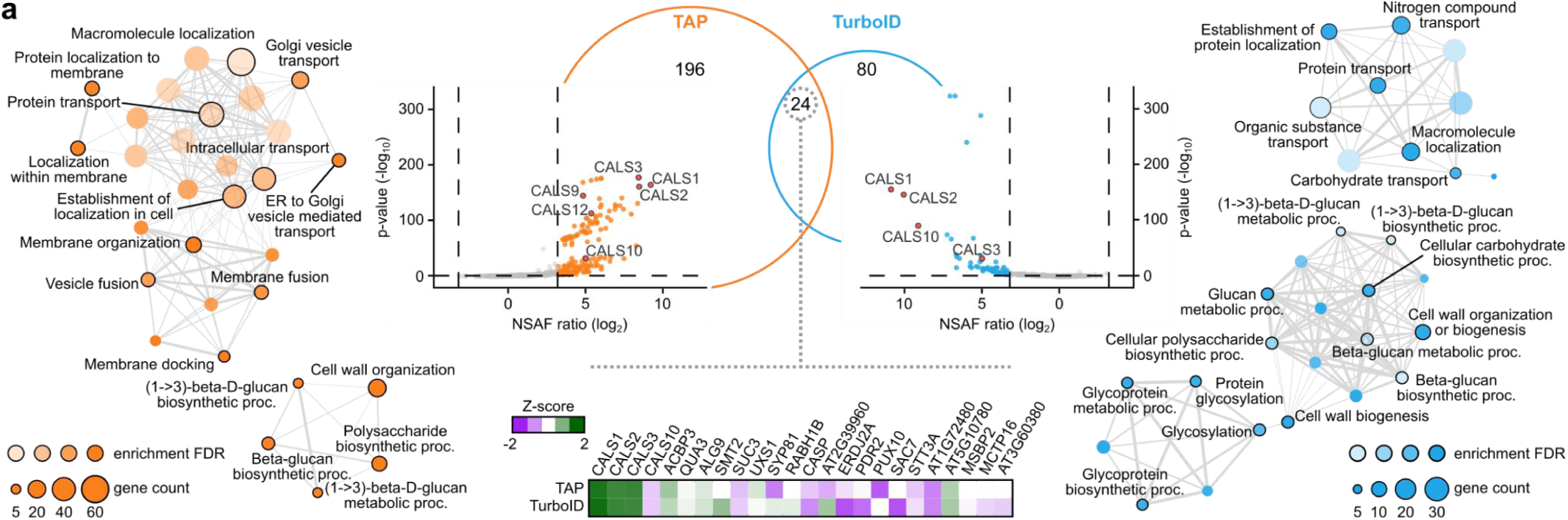
Protein interactome of CALS1. CALS1 tagged for affinity purification (AP) or proximity biotinylation (TurboID) identified putative CALS complex components (top center). The scatter plots show the significantly enriched hits, which were determined based on the protein NSAF ratio and p-value. Pink dots indicate other CALS homologs present in the dataset. Between all overlapping proteins, CALS1, CALS2 and CALS3 exhibit the highest enrichment (bottom center). The hierarchically clustered networks show the enriched GO biological process terms of the CALS1 interacting proteins present within both datasets (left, right). Each node represents a significantly enriched GO term (FDR < 0.05). The node size corresponds to the protein set size. The color hue is proportional to the FDR of each cluster. The network connectivity demonstrates the shared proteins between the individual nodes: two nodes are connected when they share at least 20% proteins, with the edge thickness depicting the number of mutually shared proteins. Selected GO terms are distinguished by a black outline.

We employed gene ontology (GO) biological process term enrichment to determine the biological function of the identified interactors. In both datasets, the most enriched terms were linked to callose synthesis, including “β-(1,3)-D-glucan biosynthetic process” and “CW organization”, together forming a distinct cluster (**Fig. 1a**). Another cluster was composed of transport-related GO terms such as “Protein transport”, “Nitrogen compound transport” or “Carbohydrate transport”. For AP-MS, this transport cluster was closely associated with membrane fusion GO terms like “membrane docking” or “vesicle fusion”. Conversely the TurboID approach revealed hits coupled with unique terms like “GDP-fucose transmembrane transport”, “Regulation of cell shape” as well as a standalone cluster of terms affiliated with macromolecule glycosylation (**Fig. 1a**).

As a complement to the GO analysis, to annotate the inter-protein relationships among the different proteins, we subjected the CALS1 interactome to the StringDB tool^30^. Additionally, a predicted subcellular localization for all hits was obtained from the SUBA5 database^31^ and used to cluster the StringDB network. The resulting network showed that the interactors of CALS1 are mostly affiliated with three main cellular compartments - endoplasmic reticulum (ER), Golgi and PM (**Fig. S1a**). A substantial number of proteins also belonged to nucleus+cytoplasm and vacuole. The majority of the 1838 interactions found by StringDB, follows the pathway of nucleus-ER-Golgi-PM. In accordance with this, all but one hit, identified by both AP-MS and TurboID, are members of the three most occupied compartments - ER, Golgi and PM, with the remaining protein being extracellular. Altogether, the analysis of our dataset by GO term enrichment, SUBA5 and StringDB databases show an intricate network of CALS1 regulators during its lifetime from proteosynthesis to its activity at the PM.

To confirm putative CALS1 interactors, we employed the split-ubiquitin assay (SUS). We selected CALS1, QSK1, a known PD protein^32^ and SMT2, a regulator of cytokinesis^33,34^, representing proteins active in canonical callose-dependent processes. In our approach, we used CALS1 as well as membrane-inserted QSK1 or SMT2 as bait, while CALS1 was utilized as the prey. Among the screened proteins, there was a strong interaction between Nub-CALS1 and Cub-CALS1. Additionally, we observed weaker interactions between CALS1 and QSK1 and SMT2 (**Fig. S2a**). These results further reinforce the CALS1 interactome dataset, validating the individual proteins’ association with CALS1 under different conditions, in both native and heterologous system. In summary, the combined data from AP-MS, TurboID and SUS corroborate the CALS1 interactome and provide a basis for future studies of callose production as regulated by various CALSC components.

### CALSC subunits mutually regulate plasmodesmal callose deposition

Even though the ability of CALS to form higher order assemblies was proposed before^10,17,18,21,22,35^, a precise composition of different CALSCs and their role in plant development has not been described. Therefore we further investigated the most enriched CALS1 homologs in our dataset, CALS1 and CALS3. Full length CALS1 (mEGFP-AtCALS1_FL_), expressed heterologously in tobacco BY-2 cell culture cells, localized to the PM and the nascent cell plate (**Fig. 2b**). Although this is consistent with earlier published data^2^, we also observed a distinct punctate pattern of mEGFP-CALS1_FL_ at anticlinal and periclinal cell walls (**Fig. 2b**). A similar pattern was obtained after aniline blue staining, which labels callose, supporting the PD identity of the observed spots (**Fig. 2a**). A small portion of the GFP signal originated from dot-like structures in the cytoplasm and at the nucleus periphery, reflecting the CALS1 population undergoing trafficking from ER through Golgi and further to PM (**Fig. 2b**).

**Fig. 2:**
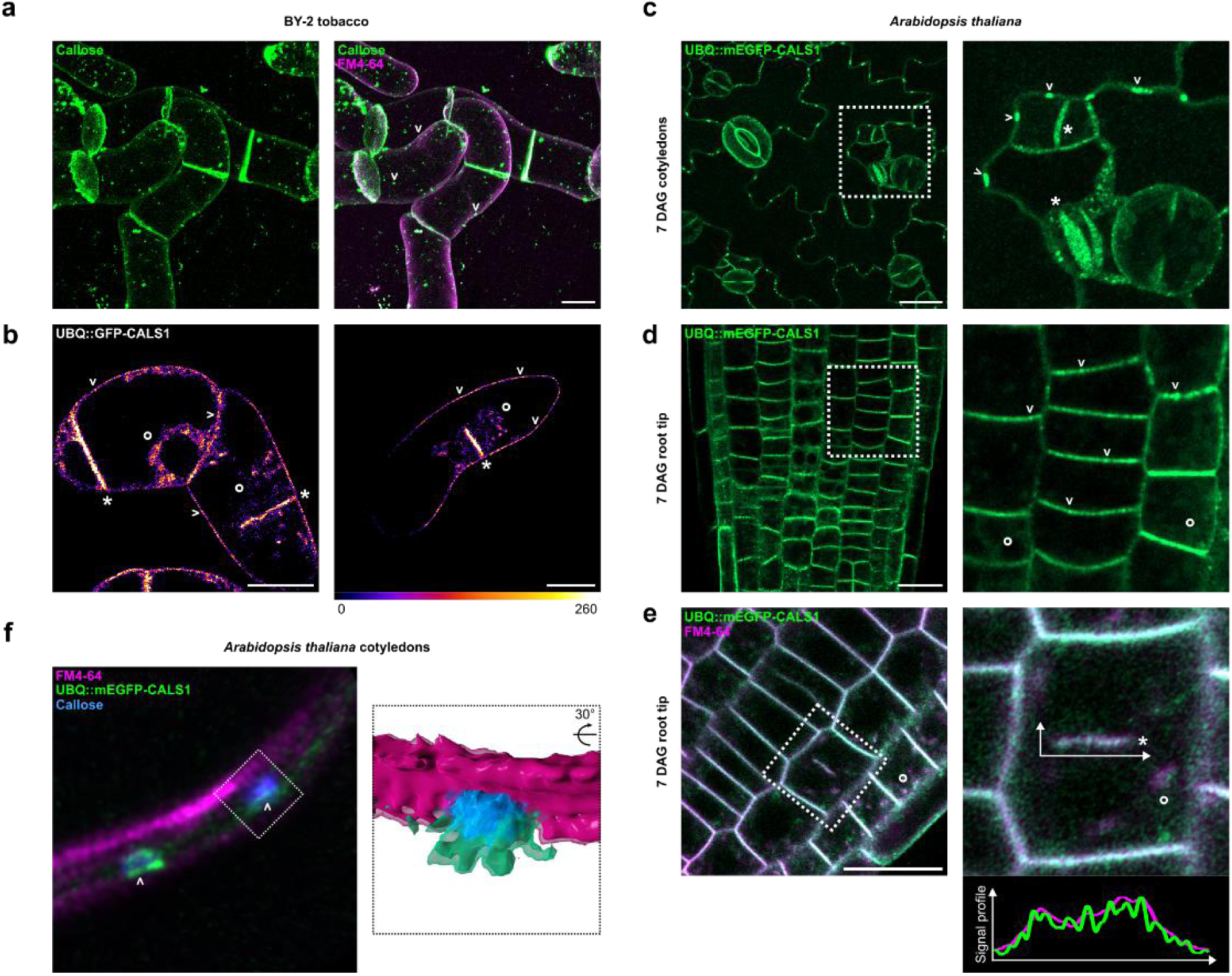
CALS1 subcellular localization *in planta*. (**a**) WT BY-2 tobacco cell culture, stained with aniline blue (green) and FM4-64 (magenta), maximum intensity projection. (**b**) 3 day-old BY-2 tobacco cell culture expressing mEGFP-CALS1 (Fire LUT). (**c**) Cotyledons of *Arabidopsis thaliana* seedlings 7 days after germination (DAG), stably transformed with mEGFP-CALS1 (green). (**d**) 7 DAG Arabidopsis root expressing mEGFP-CALS1 (green). (**e**) 7 DAG Arabidopsis root expressing mEGFP-CALS1 and counterstained with FM4-64 membrane dye (magenta). The signal profile within the growing cell plate is indicated. (**f**) Cotyledons of *Arabidopsis thaliana* seedlings 7 days after germination (DAG), stably transformed with mEGFP-CALS1 (green) and co-stained with aniline blue (magenta) and FM4-64 (magenta). Volumetric representation of the highlighted structure is provided to the right. The cell plate (*), plasmodesmata (v) and endomembrane system (°) are indicated. Scale bar = 20 μm.

To study the localization of AtCALS1 in a non-heterologous system, we generated *pUBQ-mEGFP-AtCALS1_FL_* expressing *Arabidopsis* lines. Similar to our BY-2 observations, CALS1 marked the PM with a strong enrichment at PD (**Fig. 2c,f**). A low amount of the GFP signal was also apparent in dot-like structures in the cytoplasm of root cells, likely representing transport vesicles containing CALS1 (**Fig. 2d**). Furthermore, we were able to detect punctate CALS1 signals at the growing cell plate of the dividing root tip meristem cells and cotyledon pavement cells (**Fig. 2e**), where it likely decorated the emerging primary PD^36^. We did not detect the pUBQ-mEGFP-AtCALS1_FL_ signal at places of cotyledon cells wounding, pollen tube callose plug formation or trichome-associated callose Ortmannian ring growth, suggesting that CALS1 is specific for the production of PD-associated callose (**Fig S3**). This confirms earlier *cals1* mutant analyses that suggested its role in PD regulation^37,38^.

Similar to the CALS1, multiple studies have linked the second most enriched interactome homolog, CALS3, with PD callose deposition^4,39^. In *Arabidopsis*, CALS reportedly underwent recent gene duplications^40^, forming the CALS1-CALS4 clade and possibly underlying functional redundancy between the homologs. It is likely that this evolutionary and functional relatedness underlies the absence of easily observable mutant phenotypes under normal conditions^1,41,42^, making the analysis of *cals* single mutants rather limiting. Therefore, in addition to the already characterized *cals1* and *cals3* single mutants^4,41^, we examined a *cals1/cals3* double mutant line. In particular, we checked whether they exhibit an impaired callose deposition at PD pit fields. To uncover PD pit fields without usage of classical PD protein markers such as PDLP, which are known to promote callose accumulation^43,44^, we used negative PD staining by PI^45^. The *cals1/cals3* double mutant, but not the *cals1* or *cals3* single mutants, exhibited a marked reduction in PD callose deposition in cotyledon epidermal cells (**Fig. 3a**), demonstrating that both genes are redundant for callose accumulation at PD. Interestingly, we observed an increased diameter of plasmodesmata pit fields in *cals1*, *cals3* and *cals1/3* lines compared to WT, suggesting an interplay between callose deposition and plasmodesmata development (**Fig. 3a**). Subsequently, by tracking the GFP spreading of GFP in epidermal cells of *cals1/cals3* we analyzed the effect of the impaired PD callose production on the intercellular transport. Functional assays revealed that the single mutants did not differ in basal connectivity, while exhibiting impaired closure in response to SA. In contrast, the *cals1*/*cals3* line showed an increased connectivity both in control conditions and upon SA treatment (**Fig. 3b**), further supporting the collaborative action between CALS1 and CALS3 in PD callose deposition. Complementing this, we investigated the callose patterns in the roots of the mutant lines. The major observed differences were linked to the callose deposition in the primary PDs at the cell cross-walls. While in WT the callose is visible in the matured center of the cross wall, approximately half of the *cals1* and *cals3* single mutant cross walls feature a donut-shaped structure, characterized by a lack of callose signal at its center (**Fig. 3c**). The number of donut-shaped cross walls was even higher in the *cals1*/*cals3* double mutant (**Fig. 3c**). Despite this, no defects in cytokinesis were observed in any of the mutant backgrounds, indicating that the reduced callose levels specifically affect PD without impairing callose deposition during cell plate formation. In addition, these results imply the tight regulation of the CALS activity, with processes in close spatial proximity seemingly mediated by different CALS homologs^40^.

**Fig. 3:**
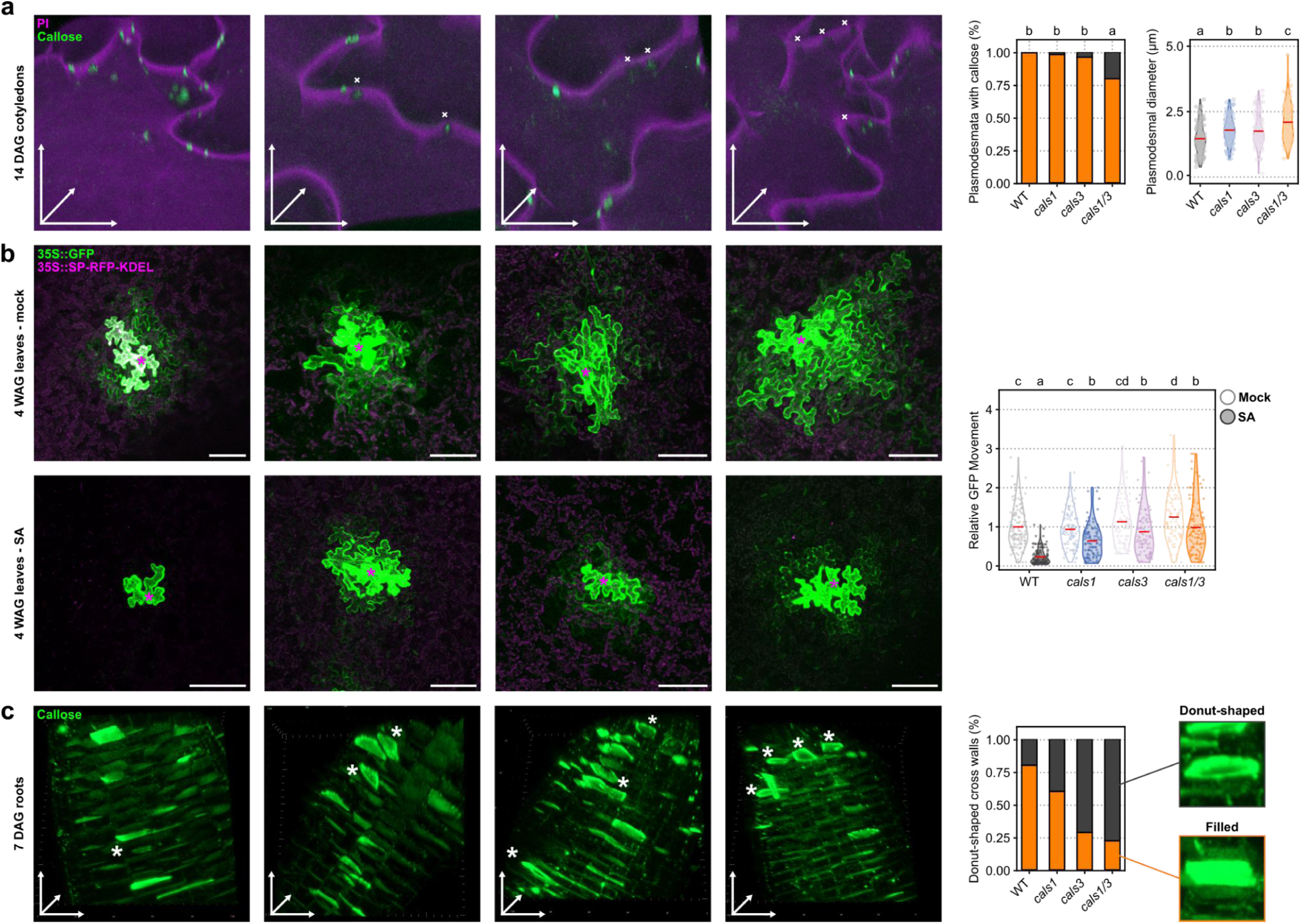
Phenotypic analysis of *cals1*, *cals3* and *cals1/3* mutant lines. (**a**) Epidermal cells of 14 (DAG) Arabidopsis cotyledons, stained with propidium iodide (PI) (magenta) and aniline blue (green). The crosses indicate plasmodesmata (PD) pit fields without a characteristic callose spot. (**b**) Plasmodesmata permeability analysis in *Arabidopsis thaliana*. Leaves bombarded with free GFP (green) and ER marker (magenta) constructs were treated with salicylic acid (SA), stimulating plasmodesmal closure. The data were normalized to the WT x mock group (**c**) 7 DAG Arabidopsis roots stained with aniline blue (green). Newly-formed cross walls with affected callose deposition in the primary plasmodesmata, demonstrated as "donut" shaped cross walls, are marked with an asterisk. Phenotype quantification of the traits is always shown to the right; the red line indicates the mean, statistically significant groups are denoted by letters, determined by one-way ANOVA with Tukey post hoc. Scale bar = 20 μm.

The *in planta* experiments provide compelling evidence that CALSC, containing CALS1 and CALS3 subunits, functions specifically at PD. We therefore compared the AP-MS and TurboID datasets with existing Arabidopsis PD proteomic datasets^46,47^. While we did not employ any subcellular compartment enrichment steps while isolating the CALSC binding partners, a non-negligible part of the interactome has been described as bona fide PD proteins (**Fig. S2b**). A total of 56 hits overlapped with the Fernandez-Calvino dataset^47^. This subset contained multiple SNARE complex members, including three SYP proteins, NSF and ASNAP2 as well as the synaptotagmin SYT5, previously shown to anchor the ER at the PM contact sites^48^. Several PM-located ion and small molecule transporters were also enriched in this subset (**Fig. S2b**). On the other hand, only two proteins were shared between the CALS1 interactome and the Brault dataset^46^. The mutual overlap of all datasets showed 13 proteins, out of which 4 were CALSs, including CALS1-3 (**Fig. S2b**). Among the overlapping proteins, 3 representatives from the MCTP multigene family, known regulators of symplastic transport^46,49^, were present. Interestingly, MCTP localization is controlled by the action of the recently characterized PI4P phosphatase SAC7^50^, another CALS1 interactor. The enrichment of MCTPs within the CALS1 dataset could hint at their collaborative action at PDs. Curiously, the CALS1 interactor QSK1 kinase, while absent in the PD datasets, also exhibits a stress-dependent relocalization to PD, affecting the callose production^32^.

Altogether our data strongly supports the role of the CALS1 and CALS3-containing CALSC governing cell-to-cell transport during plant development. While our experiments on *cals1/cals3* lines showed a marked reduction in PD-associated callose, the aniline blue signal was still not fully abolished. We believe this is caused by the residual activity of additional CALS homologs, namely CALS2 and CALS4. Indeed, together with the CALS1 and CALS3, CALS2 has been shown to respond to the *Phytophthora brassicae* effector RxLR3, inhibiting callose production and promoting PD transport^38^. Altogether, these results provide a foundation for future analyses of the relationship between CALSC composition and the production of callose during distinct cell processes.

### Core callose synthase complex adapts a trimeric architecture

The plethora of CALS1 interactors identified by our approach gives an unprecedented insight into the intricate CALSC network. Despite this, it provides only little information about the molecular architecture of the plant CALSC. We therefore decided to investigate the structural arrangement of the CALSC and its subunits. First, to determine the size of the catalytically active CALSC core, we isolated the PM fraction from the stable BY-2 tobacco line expressing *pUBQ10::mEGFP-CALS1*. This fraction, confirmed to be enzymatically active (**Fig. 5a**), was subsequently separated by blue native (BN)-PAGE. The enriched CALSC exhibited a strong enrichment around 750 kDa (**Fig. 5a, Fig S5b**). This is in line with previous studies showing the size of a glucan-synthesizing complex at roughly 750-800 kDa^10,22,35^, which corresponds to the CALSC core consisting of three CALS subunits. Additional bands at 250 and 500 kDa possibly reflect the different CALSC stoichiometries in the sample, perhaps linked to the stepwise assembly of the CALSC.

We then utilized crosslinking (XL)-MS to analyze the spatial arrangement of the CALSC components. Therefore, CALS1-GS^rhino^ was purified from *Arabidopsis* culture cells, followed by on-bead crosslinking by Bissulfosuccinimidyl suberate (BS3). We tested BS3 concentrations of 0.6, 1.2, 2.4 and 4.8 mM, out of which even the lowest concentration was sufficient to crosslink the complex (**Fig. S5b**). Among the proteins identified by pFind in the obtained MS spectra were multiple CALS homologs, including CALS1-3, which were the top 3 hits. Using pLink^51^, we generated a dataset that contained 13 unique crosslinked sites between either CALS1, CALS2 and CALS3, and CALS10. Moreover, additional 13 sites between Streptavidin and CALS1, CALS2, CALS3 or CALS4 were determined. For the rest of the proteins identified by XL-MS (HSP70-1, ACC2 and EF1) the analysis found no significant crosslinks (XLs) with CALS1 (**Supplementary data**).

Although the analysis discovered multiple XLs between CALS1, CALS2 and CALS3, their high sequence similarity^40^ did not allow to pinpoint individual crosslinked peptides to a specific homolog and thus enabled only to consider them as intramolecular. Since there is no experimental structure of plant CALS solved to date, we relied on protein structure prediction by AlphaFold2 (AF2) to generate the input model. Recently, there has been a surge of studies showing that AF2 predictions of ScFKS1 are highly confident and almost identical to the experimental models obtained by cryo-EM^13–15^. Indeed the AF2 prediction of CALS1-3 monomers yielded high-confidence models (**Fig. 4b, Fig. S4a**). In addition, we superimposed the predicted CALS structure with the published cryo-EM structure of yeast FKS1^14^ (**Fig. S4c**). The RMSD below 5Å suggests a close similarity between the fitted structures, reflecting the evolutionary relatedness of these two glucan synthases^40^. When the XLs were mapped to the AF2 models of either CALS1, CALS2 or CALS3 as strictly intramolecular, 92% satisfied the 30Å Cɑ-Cɑ distance^52^ (**Fig. 4a**). Only a single XL type, between the residues K302-K453, both located at a flexible region, violated this cutoff limit. These results strongly support the overall reliability of the monomer structure model.

**Fig. 4:**
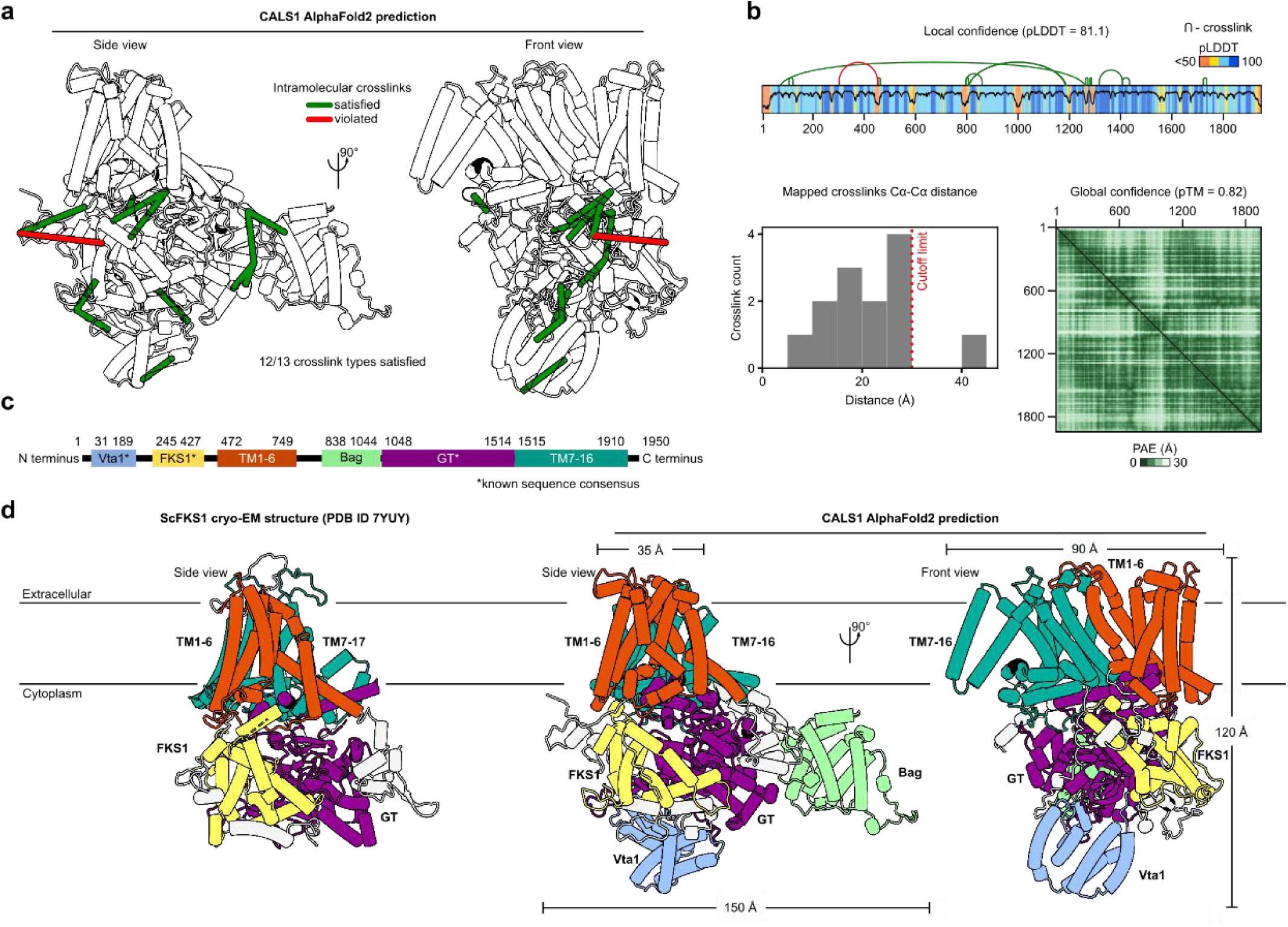
Mapping crosslinked peptides on the CALS1 AlphaFold2 (AF2) structural model. (**a**) Ribbon representation of the CALS1 with BS3 pseudobond crosslinks (XLs) mapped to the structure. The crosslinks’ distance was measured between the Cαs of the crosslinked residues. To distinguish the satisfied (green) and violated (red) XLs, a cutoff of 30 Å was applied. (**b**) Histogram of the detected XLs, with the cutoff limit indicated. The structural model exhibits a high local confidence pLDDT score (blue = high confidence, orange = low confidence). The overall model quality is validated by the predicted alignment error (PAE) plot. (**c**) CALS1 functional domains by InterPro aligned to the CALS1 sequence. Domains with a previously known sequence consensus are depicted (**\***). (**d**) Structural comparison of CALS1 and its budding yeast homolog ScFKS1 (PDB 7YUY). The general architecture, catalytic core and the transmembrane (TM) helix positions between the proteins is maintained. In CALS1, there are two additional cytoplasmic domains, the Vta1 (pale blue) domain and the Bag (pale green) domain.

**Fig. 5:**
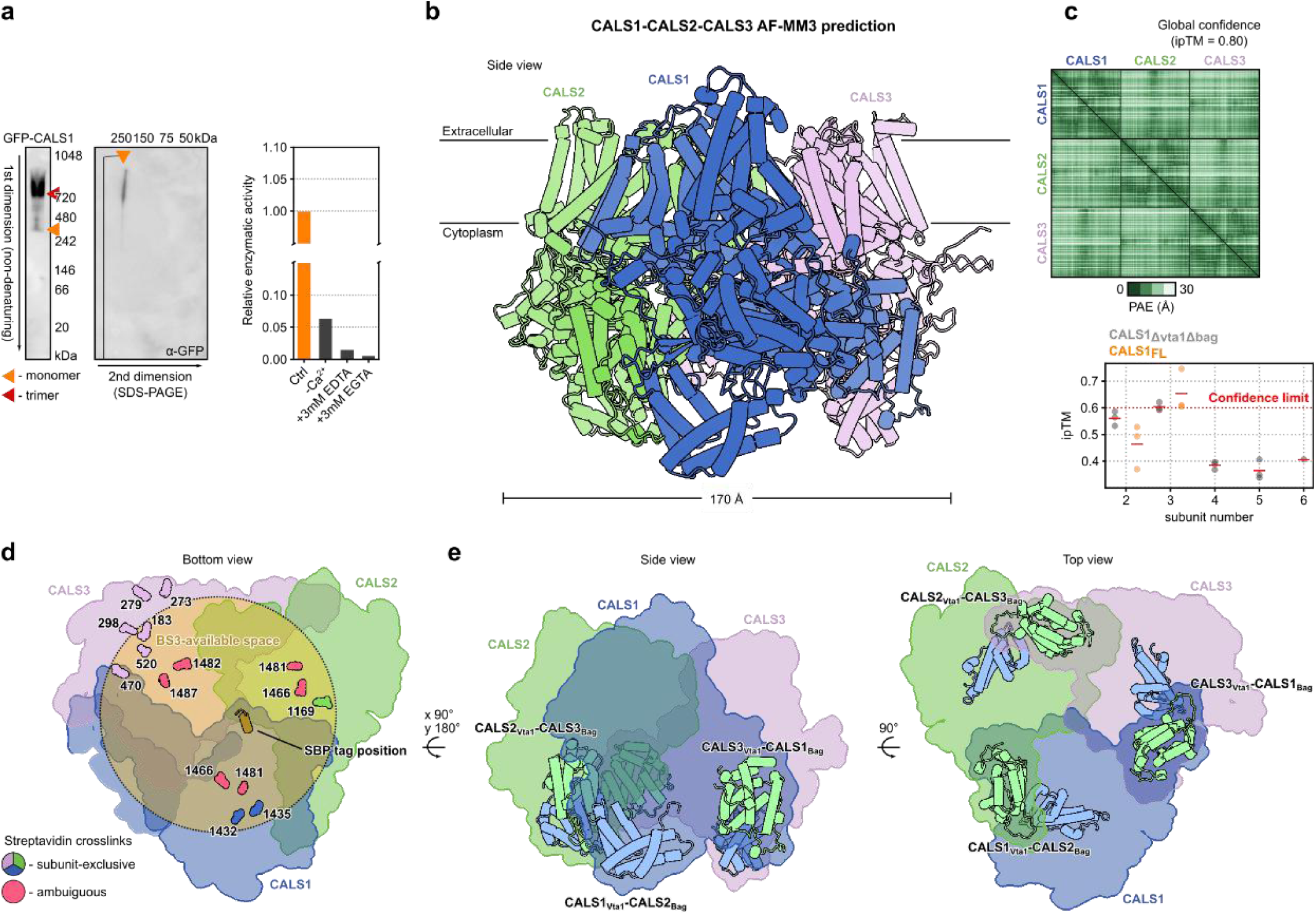
Overview of the CALS complex (CALSC) core architecture. (**a**) Blue native PAGE of a plasma membrane fraction from BY-2 cells expressing GFP-CALS1. The size of the strongest band corresponds to the trimeric constitution of the CALSC. Enzymatic activity of the fraction under different buffer conditions is shown to the right. (**b**) AF2-MM3 model of the CALS1-3 heterotrimer. The presumed integral membrane parts are positioned within a common region, while the cytoplasmic regions cluster to a single large domain. The global model quality is validated by a low predicted alignment error (PAE). (**c**) AF2-MM3 predictions of CALSC stoichiometries containing 2-6 subunits of CALS1 or its truncated version. Based on the mean ipTM score of the top three models, the trimer is the most trustworthy assembly. (**d**) Streptavidin-CALSC BS3 crosslinks mapped to the CALSC heterotrimer. The SBP tag at the C-terminus of the bait CALS1 is indicated by a brown helix. Space permeable by BS3 is depicted by an orange circle. The crosslinked residues are indicated by a van der Waals representation. (**e**) Position of the Vta1 and Bag domains within the CALSC trimer. The domains lie at the interface between the individual subunits.

In a similar manner, AF2-multimer3 (AF2-MM3) was employed to predict the structure of multiple-CALS subunit CALSCs. We tested the assembly of 2-6 subunits, out of which only the trimeric assembly showed a confident iPTM score of over 0.6^53^ (**Fig. 5b**). To analyze the preference for certain homologs, we produced both homo-(containing only CALS1 subunits) and heterotrimeric (composed of CALS1, CALS2 and CALS3) CALSC assemblies (**Fig. 5c**). We note that there was no substantial difference between the confidence scores of the homo- and heterotrimers (**Supplementary data**). In connection with our interactome data we thus chose the CALS1-CALS2-CALS3 heterotrimer (coined as the CALSC trimer) for all further analyses.

Because the XLs we obtained could not be utilized unambiguously to analyze the intermolecular linkages, we considered the Streptavidin XLs as an indirect way of validating the CALSC model. In the last step of the TAP, Streptavidin-coated beads bind the SBP tag at the C-terminus of the CALS1 bait. The presence of multiple XLs between the Streptavidin and a specific CALS residue (including CALS2 and CALS3 prey) implies their close spatial proximity. Indeed upon mapping to the CALSC trimer, the residues crosslinked with Streptavidin were largely restricted to the cytoplasmic side of the protein, inside the cavity at the interface of the three homologs. This is in line with the orientation of the SBP within the same cavity (**Fig. 5d**). On top of this, the majority of the crosslinked residues fell within the 30Å cutoff distance, which together with the BN-PAGE results and the AF2 scores further validates the trimeric assembly hypothesis.

### Arabidopsis CALS structure reveals two distinct plant specific domains, which are key elements of the CALSC subunit interaction

The general CALS topology includes two cytoplasmic regions, located at the N-terminus and in the middle of the protein. CALS contains 16 TM helices – TM1-6 separating the cytoplasmic regions and TM7-16, positioned between the central cytoplasmic region and a short C-terminus. The overall CALS topology is similar to the ScFKS1 experimental models – the hydrophobic residue-rich TM helices assemble into a single integral membrane region, out of which both cytoplasmic regions protrude (**Fig. S3e**). In CALS1, several functional domains have been identified, including the Vta1, FKS1 (both within the N-terminal region) and the GT domain (making up the bulk of the central cytoplasmic region) (**Fig. 4c,d, Fig. S6a**). By superimposing CALS1 and FKS1, we noticed several distinct differences. The helical Vta1 domain was absent in the yeast counterpart and an additional well-folded globular part was present in CALS1 (**Fig. 4d, Fig. S4c**). Even though the latter belongs to the large cytoplasmic region, its position is independent from the catalytic GT domain, extending outside the core of the protein. Further attempts to classify this region by the DALI and Foldseek structural similarity search tools^54,55^ yielded no related experimental structures and no structurally similar proteins outside of the CALS family. We, therefore, hypothesized that this region, which we titled the Bag domain, is likely a plant specific addition to CALS. To test this hypothesis, we predicted the monomeric structures of CALS orthologs across the plant lineage and fungi. The conserved FKS1 and GT domains were present in all tested species. On the other hand, Vta1 and Bag domains were limited to the plant orthologs, including partial Vta1-like folds occurring in chlorophyte algae (**Supplementary data)**. Evolutionary conservation analysis by ConSurf revealed that both Vta1 and Bag domain are only mildly conserved compared to the rest of the protein (**Fig. S4d, Fig. S6b**). This strongly suggests that Vta1 and Bag domains were newly acquired by the plant-specific CALSs, with the former undergoing a partial secondary loss in chlorophyte algae (**Fig. S6c**).

Within the CALS monomer both the Vta1 and Bag domain are positioned away from the central core of the protein, facing the cytoplasm (**Fig. 4d**). Contrarily in the model of the CALSC trimer, the plant specific domains are at the interface between the respective subunits (**Fig. 5d**). The buried area between the individual subunits is between 2730-3020 Å^2^ (**Fig. S5c**), out of which the Bag-included interface accounts for more than a half (**Supplementary data**). We hypothesized that the presence of the Bag domain possibly stabilizes the CALSC trimer. To test this, we performed SUS between the full length CALS1 (CALS1_FL_) and the CALS1 Bag domain (CALS1_Bag_) as the Nub-tagged prey. Compared to the binary interaction between CALS1_FL_ as both bait and prey, CALS1_Bag_-expressing cells exhibited comparable colony growth, validating their mutual interaction (**Fig. S1a**). On the other hand, the interaction was fully abolished for CALS1 lacking the Bag domain (CALS1_Δbag_).

Next, we investigated the association between mEGFP-CALS1_FL_ and mScarlet-CALS1_Bag_ in *Nicotiana benthamiana*. mEGFP-CALS1 localized predominantly to the PM, with strong enrichment in distinct PD-like PM domains, while mEGFP-Bag domain expressed by itself was mostly cytoplasmic due to its lack of membrane-binding elements. Despite this, upon coexpression with mEGFP-CALS1_FL_, a small mScarlet-Bag domain population was enriched in PM domains and was also present in the microsomal fraction (**Fig. S1b**, **Fig. S5d**), implying that the Bag domain itself might interact with PM-resident proteins, including CALS..

The Vta1 domain was likewise present in the microsomal fraction, albeit to a lesser extent compared to the Bag domain (**Fig. S5d**). These differences might reflect the different capacity of the plant-specific domains to mediate the CALSC oligomerization. On the other hand, the cytoplasmic orientation of the Vta1 and Bag domains in the monomeric CALS could be also relevant for their interaction with non-CALS proteins, thereby influencing the equilibrium between the assembled and disassembled complexes and possibly CALS activity or compartmentalization. An analogous mechanism of oligomerization has been identified in CESA, mediated by the trimer-stabilizing plant-specific P-CR domains, effectively promoting the crystalline ultrastructure of cellulose microfibrils^56–58^. It has been shown that FKS1 oligomerization through a FKS1-tRNA-FKS1 complex inhibits its enzymatic activity^59^. The mode of interaction within such structure differs markedly from the CALSC trimer depicted here; this could be due to the absence of the Vta1 and Bag domains in the yeast glucan synthase. As such, further investigation into the plant specific domains and their role in CALSC will be key for understanding the biological relevance of the CALSC oligomerization.

### Evolutionary conserved motifs control the enzymatic activity of CALS1

One of the key structural elements of CALS1, located at the central cytoplasmic part, is the GT domain, (**Fig. 4d**). The GT domain is hypothesized to contain the catalytic pocket (**Fig. 6a**) that mediates the transfer of the glucose unit from UPG to the nascent glucan chain^10,17,60^, in which multiple extremely conserved amino acid motifs (**Fig. 6b**) have been identified, including the SED1, SED2, RxTG and NQD motifs^60,61^. Within the GT domain, all motifs are positioned in close spatial proximity to each other (**Fig. 6a**). CALS belongs to the GT48 family of inverting glycosyltransferases and as such catalyzes the glycosidic bond formation between the C1 atom of the UPG donor D-glucose and the C3 atom of the nascent glucan^62^. The formation of the glucan chain requires additional ligands that coordinate the reaction, including Ca^2+^ and Mg^2+^ ions^15,21,24–26,59^. To this date, the spatial interplay of the substrate, product and both ligands within glucan synthase active site remains to be characterized. To investigate the topology of the known reaction components we generated an AlphaFold3 model containing the UPG, Ca^2+^ and Mg^2+^ and the product laminaribiose (β-D-glucose-1,3-β-D-glucose). The predicted components assumed position within the putative pocket, with the pLDDT and PAE confidence scores showing highly confident topology (**Fig. 6a,c,d**). In detail, the UPG molecule was located at the entrance to the pocket, its pyrophosphate group proximal to the C3 of the laminaribiose was predicted to be encased within the pocket. Their mutual interaction was coordinated by Ca^2+^ and Mg^2+^ positioned in between UPG and laminaribiose. The orientation of the molecules was in accordance with the reaction progression - the UPG cleavage provides energy for the glycosidic bond formation between the acceptor glucan and the donor glycosyl molecule, while the coordinating metal ions mediate the electron transfer^62^.

**Fig. 6:**
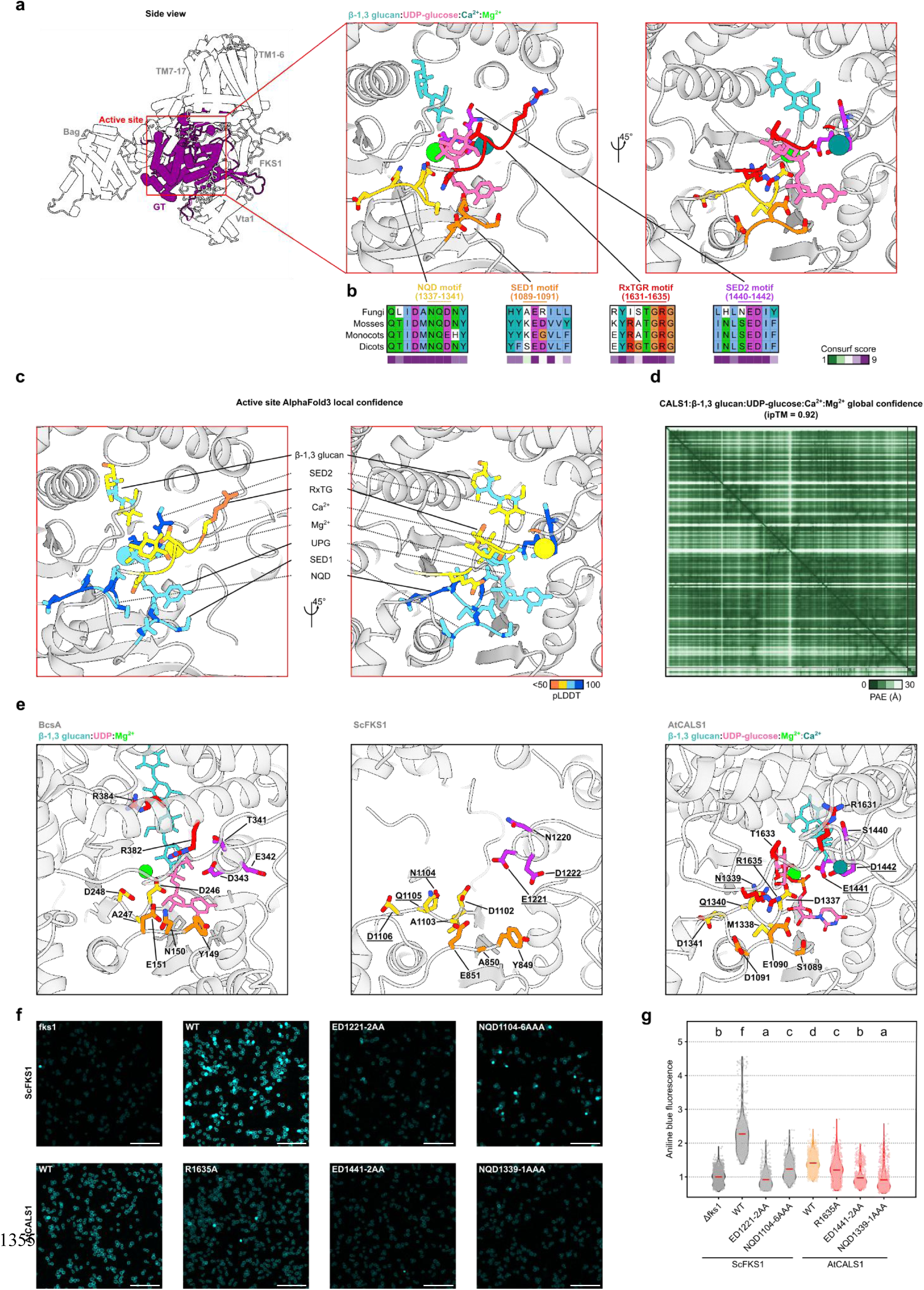
CALS1 active site overview. (**a**) Location of the active site within the full structure. (**b**) Conserved active site motifs mapped on the CALS1 structure including the reaction components. The multiple sequence alignment of the individual motifs and ConSurf score highlight the evolutionary conservation of active site residues. (**c-d**) AlphaFold3 metrics of the prediction displayed in **a**. A confident score was achieved for all components and their coordinating residues. PAE plot results favor the global arrangement of the predicted molecules. (**e**) Spatial superimposition of active sites of CALS1, yeast FKS1 and bacterial CESA. The evolutionary conserved residues in atomic representation are colored uniformly by the common motif. The underlined residues signify amino acids selected for the yeast glucan content assay. (**f**) Glucan content assay of yeast Δfks1 complementant lines. (**g**) Quantification of the glucan content measured in **f**. The aniline blue fluorescence of each line was normalized to the Δfks1. The red line depicts the mean value, data were analyzed by the Kruskal-Wallis test with Dunn post hoc analysis. Scale bar = 20 μm.

To validate the AF3 prediction, we further investigated the amino acid moieties of the catalytic pocket. Within the model, the UPG binding was facilitated by a short RxTGR motif loop, the particular interaction occurring between the positively charged R1635 side chain and the negatively charged UPG pyrophosphate (**Fig. 6e**). The RxTGR loop lies opposite to the SED1, from which E1090 coordinated the uridine moiety of UPG (**Fig. 6e**). Although in yeast the full SED1 motif is not conserved, the structurally homologous residues Y849 and E851, along with K1082 and N1085, perform a similar role^14^. This topology is maintained also in the bacterial CESA which partially shares its active site fold with β-1-3 glucan synthases (**Fig. 6e**). More specifically, the residues Y149, E151, K226 and K229 are binding uridine, while the RxTGR motif is not present and the pyrophosphate group is coordinated by R382 (a part of an RGR motif) instead^63^ (**Fig. 6e**). Next, we examined the coordinating metal ions within the structure. The binding of the Mg^2+^ was mediated mutually by the NQD motif residue D1337 and the UPG pyrophosphate (**Fig. 6e**), similar to the bacterial CESA residue D236^63,64^. In contrast, a recent FKS1 study has proposed that residues H1198, D1197 and C1244, located approximately 8 Å away, bind the Mg^2+^; this was linked to the lack of the UPG within the pocket, their mutual interaction promoting the correct progress of the reaction^59^. To go further, in our model the Ca^2+^ ion was positioned next to the E1441 and D1442 of the SED2 motif (**Fig. 6e**). Previous studies have implied that the relevant yeast homologous residues E1221 and D1222 stabilize the nascent glucan^15^ or act as a catalytic base^63^. Their mutation led to severe inhibition of the enzymatic activity, although no connection to the calcium binding has been made. Nevertheless, further investigation of the relationship between metal ion cofactors and glucan synthase active site residues is required to determine their precise role during glucan synthesis.

We then decided to test the role of the active site residues in regulating the CALS activity *in vivo*. Multiple studies have shown that the activity of fungal glucose synthases can be abolished by generating point mutations in the protein^13–15,59^. Moreover, the ability of plant CALSs to rescue yeast *fks1* mutants has been demonstrated before^61^. In our experimental setup, we introduced pTDH3::6xHA-CALS1 mutated in selected active site residues into the BY21271 yeast strain (Δfks1, **Supplementary data**) deficient in glucan synthesis^65^. Subsequently we stained the complementation strains with aniline blue and analyzed CW glucan levels using confocal microscopy. As a control, we utilized the WT FKS1 and mutant variants in the essential^13–15^ NQD (residues 1104-1106) and ED2 (residues 1221-1222) motifs. The expression of the complementing cassette was confirmed by WB (**Fig S8a**).

Compared to the Δfks1 strain, yeast expressing CALS1_WT_ exhibited an increased aniline blue signal (**Fig. 6f,g**). This increase (approximately 2.5x higher) was even more striking in the ScFKS1 overexpression lines. In CALS1_NQD1339-1AAA_ and CALS1_ED1441-2AA_, the glucan content was similar to the levels in Δfks1; in the former it was even lower than the negative control, which might be attributed to a reported compensatory effect of ScFKS2 and FKS3^66,67^. As in the previous works^14,15^, the FKS1_NQD1104-6AAA_ and FKS1_ED1221-2AA_ mutants exhibited reduced activity (**Fig. 6f,g**).

In a similar way, we examined the effect of RxTGR motif manipulation on CALS1 activity. In the Δfks1 background, CALS1_R1635A_ overexpression reduced the glucan content compared to the CALS1_WT_, although it remained slightly higher than in the CALS1_NQD1339-1AAA_ and CALS1_ED1441-2AA_ (**Fig. 6f,g**). Next, we decided to analyze the effect of the mutation on the RxTGR-UPG interaction further. We generated all atom molecular dynamics (MD) systems containing CALS1_WT_ and CALS1_R1635A_ embedded in a lipid bilayer. Additionally we provided laminaribiose, UPG and both Mg^2+^ and Ca^2+^. We observed that after 500 ns simulation, no dramatic change occurred within the system. It is likely that the presence of multiple highly charged atoms within close proximity stabilizes their topology and does not allow the movement of the reactants. As an alternative, we tried to simulate the endpoint state of the reaction catalyzed by CALS, containing only 3xBGL and UDP. In the WT setup, the UDP remained bound to R1635 throughout the whole 500 ns simulation, in all 5 simulation repeats (**Fig. 7g**). Contrastingly, the CALS1_R1635A_ exhibited an increased RxTGR movement and resulted in the UDP exit in 2/5 repeats. We hypothesize that the R1635-UPG interaction stabilizes the substrate in its active site position and promotes the correct orientation of the pyrophosphate group. It can be further speculated that the RxTGR outward movement enlarges the catalytic pocket entrance, promoting the UDP for UPG exchange. Our data are consistent with previous FKS1 results, where the RxTGR-analogous loop movement has been proposed to stimulate the catalytic activity^16^. Interestingly, a similar mechanism of substrate entry regulation was demonstrated for the poplar CESA8 gating loop^68^. Altogether, the combination of the yeast glucan content and MD simulation data highlights the importance of R1635 in callose synthesis. However, the mild effect we observed in both instances could be attributed to the presence of other UPG-binding residues or to the rest of the RxTGR loop, and further studies into UPG-binding regulation are necessary. Still, these results provide unprecedented insight into the evolutionary conservation of β-1,3 glucan synthesis.

**Fig. 7:**
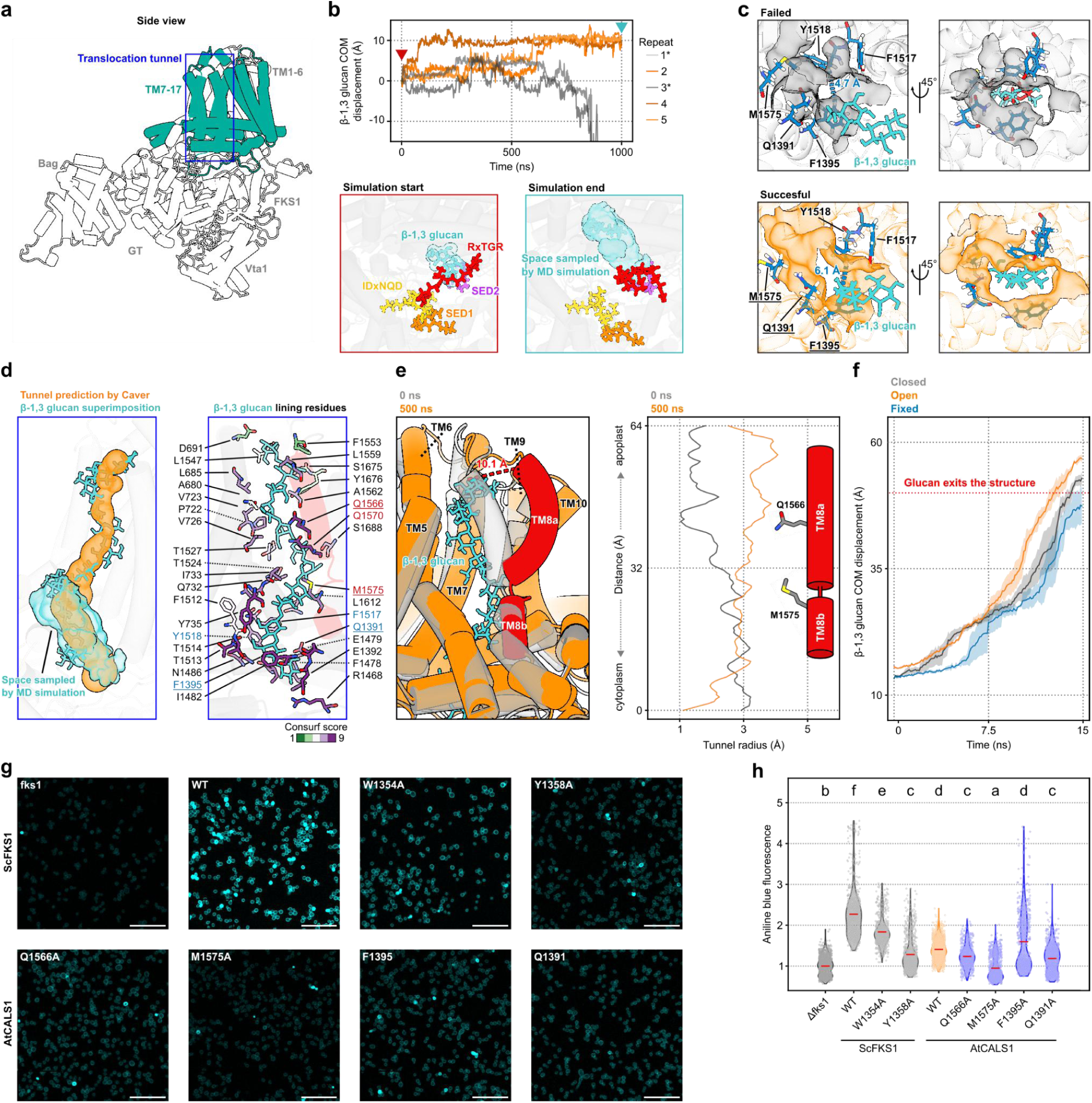
Export of the nascent glucan chain. (**a**) Location of the translocation tunnel within the full structure. (**b**) Molecular dynamics (MD) simulation of laminaribiose inside the CALS active site. Orange = trajectories with successfully transported glucan; grey = failed simulation with glucan exiting the catalytic pocket. Representative snapshots from the start and end of the simulation provided at the bottom. (**c**) Detailed view of the selectivity filter at the tunnel entrance. The topology of the glucan in relation to the filter residues is displayed for both failed and successful simulations. The blue pseudobond shows the distance between F1395 and Y1518. On the right, the glucan atoms clashing with the pocket are indicated by red color. (**d**) Translocation tunnel superimposed to glucan experimental structure and cavity sampled by MD. Tunnel lining residues are shown to the right, with residues selected for mutation analysis underlined. (**e**) Startpoint and endpoint states of the translocation tunnel containing the curdlan, simulated for 500 ns. The movement of TM8 helix (red) is indicated by a dashed pseudobond with the tunnel radius displayed to the right. (**f**) Glucan movement within the tunnel during steered MD. (**g**) Glucan content assay of yeast Δfks1 complementation lines. The glucan content was determined based on the aniline blue fluorescence using confocal microscopy. (**h**) Quantification of the glucan content measured in **g**. The aniline blue fluorescence of each line was normalized to the Δfks1. The red line depicts the mean value, data were analyzed by the Kruskal-Wallis test with Dunn post hoc analysis. Scale bar = 20 μm.

### CALS1 transmembrane helix 8 is crucial for the nascent callose chain production

While the mechanisms of the β-1,3-glucan synthesis, especially in fungi, have been described in other studies^13–16^, the molecular basis of transport of the produced glucan chain into the extracellular space still remains largely unknown. First, to determine the ability of the CALS1 structure to transport glucan and sample the putative translocation space, we performed MD simulation of CALS1 inside a lipid bilayer. The system included the laminaribiose as the model glucan, its starting position derived by AF3 (**Fig. 6a**). After 1 µs simulation time, the laminaribiose passed a distance of 10Å in 3 out of 5 repeats, translocating further into the cavity and into the integral membrane region of the protein (**Fig. 7a, b**). Curiously in the remaining 2 runs, the laminaribiose initially moved inside the catalytic pocket but eventually diffused out of the structure. These two simulation runs point at the presence of residues restricting the reverse transport of the produced glucan. Although all runs started from an identical starting point, those which were successful in transporting the glucan further exhibited a cooperation of F1395 with F1517, Y1518, Q1391 and M1575 to create a small laminaribiose-fitting pocket (**Fig. 7c**). The interaction between the glucan and F1517 moved laminaribiose deeper into the pocket, priming it for further translocation. Moreover the backwards diffusion was partially prevented by Q1391 plugging the entrance. In contrast, in the two “failed” simulations the glucan adopted a different approaching angle, unable to form a stable binding mode with the residue F1395 (**Fig. 7c**). Meanwhile the F1517 rotation away from the pocket prevented its entry. We hypothesize that the stacking interaction between the glucan ring and the aromatic side chain of both phenylalanine residues is crucial for enabling the entry of glucan at a correct angle.

To validate the *in vivo* importance of F1395 and Q1391, we generated CALS1F1395A and CALS1Q1391A mutants for glucan content measurements in yeast. While the observed signal in F1395A was statistically comparable to WT, the strain exhibited large variability, up to 4 times the Δfks1 mutant levels, suggesting that the F1395 mutation affects callose production (**Fig. 7g,h**). The Q1391A line had lower glucan content than CALS1WT, perhaps due to reduced retention of the nascent chain during callose synthesis (**Fig. 7g,h**). Comparable to our results, the mutation of FKS1 residues F1297 and H1298 (homologs of F1517 and Y1518) is lethal^14^. In the MD simulations, homologous CALS1 residues interact with the transported glucan at the far end of the active site cavity, stabilising glucan position (**Fig. 7c**).

To further identify possible pores through which glucan translocation could be mediated, we performed CaverWEB^69^ prediction of tunnels on the CALS1 structure. We filtered out the predictions ending prematurely inside the CALS structure or the lipid bilayer, yielding a tunnel delineated by the helices TM5-TM10, which connected the active site with the extracellular space (**Fig. 7d, Fig. S7d**). While previous studies on FKS1 have suggested a similar position for the transporting tunnel^14,15^, based on the superimposition with bacterial CESA, we reasoned that the different β-1,4 chemical linkage of cellulose might affect the structure of the glucan itself and promote false positive inferences. Using transmission electron microscopy, it has been shown that the ultrastructure of curdlan is comparable to β-1,3-glucan synthesized by *Arabidopsis* microsomal fraction^24,70^. To validate our findings we therefore considered the experimental curdlan structure, composed of 12 glucose subunits^71^ as the model β-1,3-glucan, which was superimposed into the putative transport tunnel, fitting into the transport cavity with only a few clashes (**Fig. 7d**). Moreover, its position was further supported by the overlap with the space sampled by the laminaribiose in the MD simulation. This position is consistent with the glucan electron density within the FKS1 structure, resolved by a recent study^16^.

One of the helices marking the transport tunnel, the TM8, has been implied as crucial for glucan deposition, providing extra space for the glucan and facilitating export^15,16^. To test this model and to investigate the dynamics of the protein during the glucan transport, we performed MD simulations of CALS1 containing the fitted curdlan. In accordance with the hypothesis, the TM8 exhibited an outward movement of approximately 10.1 Å, enlarging the tunnel radius by 3 Å (**Fig. 7e**). While the TM8 caused a substantial opening of the tunnel, the curdlan within the structure was not further translocated outside the CALS structure even after 1 µs. We hypothesized that this was likely due to two limitations of our simulation setup. Firstly, the synthesis of the callose chain requires energy supplied by the UPG cleavage, presumably necessary to accommodate the protein and “push” the nascent glucan outside. Secondly, the extracellular environment into which callose is deposited is tightly packed with other CW elements which could act as an opposing force against the nascent callose, pulling the glucan away from the protein. Because the MD limits the testing of the former hypothesis, we decided to simulate the latter by employing the steered MD simulation. As the input, we compared three distinct CALS1-curdlan setups. In negative control, CALS1 in its closed state with the TM8 fixed by position restraints was used. Alternatively, the same system without the restraints was used. Finally, CALS1 in its open state (after 1 µs run of the non-steered simulation) was tested as well. To simulate “pulling”, we applied an external force of 500 kJ.mol^-1^.nm^-2^ to the central glucose subunit. In all three simulation setups, we observed the movement of the curdlan into the extracellular space within 15 ns of computational time (**Fig. 7f**). The full translocation of the glucan occurred the fastest in the open input state, followed by the closed non-fixed state. Although the fixation of the TM8 did not prevent the translocation of the glucan, this translocation was comparably slower, enabled by the imposed artifact deformation of TM6 during the simulation (**Fig. S7d**).

To go further, we analyzed the evolutionary conservation of the residues lining the transport tunnel, confirming that several residues exhibit the highest conservation level (**Fig. S7a**). According to the ConSurf score as well as their position in contact with the curdlan, we chose two highly conserved TM8 residues, M1575 and Q1566, for glucan content measurements in yeast (**Fig. 7d**). As a control group, we tested two FKS1 TM8 residues oriented towards the central tunnel, W1354 and Y1358. Both yeast mutants exhibited lowered glucan content compared to FKS1_WT_, although it was significantly higher than in Δfks1 (**Fig. 7g,h**). The CALS1_Q1566A_ showed a similar pattern of callose production inhibition, while the CALS1_M1575A_ mutant had glucan content even lower than the Δfks1 (**Fig. 7g,h**). This is in contrast with works which show that transport tunnel residues’ mutation F1311A and F1475A leads to an increased enzymatic activity^14^. Another study has similarly pinpointed multiple residues crucial in glucan export^16^, although no direct link to the enzymatic activity has been made. While the FKS1 channel itself is lined with multiple hydrophobic residues, it is possible that the mutations we chose disrupted the TM8 structure as a whole (e.g. M1575 connects the TM8a and TM8b helices), manifested by an inhibitory effect. Despite this, our results are overall consistent with the previous structural studies of FKS1 and bacterial CESA^14–16,63^, not only confirming the necessity of the TM8 opening but also providing information about the dynamic mechanisms underlying the export of the glucan to the CW.

### Final summary of the paper

Although the localized synthesis of the β-1-3 glucan callose underlies many crucial developmental processes, previous studies have unraveled only limited information about molecular mechanisms involved and the regulatory networks controlling its deposition into the CW. Here, we identify a large portfolio of binding partners comprising the central callose-synthesizing unit, the CALSC. The core of the CALSC adopts a trimeric architecture, mediated by a mutual interaction of three CALS subunits. The enrichment of homologs CALS1 and CALS3, together with the distinct composition of the interactors, likely governs the specific activity of this CALSC at PD, as shown in our *in planta* microscopic experiments. Our investigation of *cals1/cals3* mutants provides a detailed view of the developmental and physiological consequences arising when the subset of PD cannot be closed or regulated by callose deposition.

In addition, through combined computational and experimental approaches we characterize the molecular architecture of the CALSC core. By comparison with the previously published structural models of the fungal homolog of CALS, FKS1^13,13–15^, we determine key structural elements, the Vta1 and Bag domains, positioned at the interface between the CALSC subunits. We show that the Bag domain is essential for inter-CALS interactions. The novel acquisition of these domains in plants likely led to the emergence of the trimeric CALSC within the green lineage. Compared to the single-catalytic subunit glucan synthases in yeast, this presumably enabled the adaptation of differently composed CALSCs to the plethora of callose-dependent processes currently known in plants^40^. On top of this, we characterize the catalytic pocket of the CALSC and showcase the interaction between specific amino acid residues and the reactants, in an unprecedented detail. We pinpoint the translocation tunnel within the structure, connecting the active site with the extracellular space. Our data confirm the originally proposed model^14,16^, showing the need for the movement of the TM8 for efficient glucan export. Unlike the FKS1 studies^13,13–16^, our approach expands knowledge of the dynamics of β-1,3 glucan synthesis and transport, including the reaction components, and provides evidence of its conservation across plants, fungi, and bacteria. Therefore, this work provides a strong basis for future studies of the molecular mechanisms underlying glucan production, including easy adaptation to non-plant models.

### Limitations of the study

Although the CALS1 interactome was determined using diverse biochemical and *in vitro* methods, the final dataset reflects the CALS interactome in Arabidopsis cell culture rather than in intact plant tissue. This limits the implications for the multicellular plant body under natural conditions and for the inherent differences in CALSC composition across cell types and developmental stages. Future investigation of the CALSC regulatory network would undoubtedly benefit from utilizing several plant systems. The structural models of CALS in monomer and trimer constitutions are the result of combined experimental and computational techniques. Unfortunately, the close homology among CALS1, CALS2, and CALS3 prevents unambiguous mapping of the obtained crosslinks within the model. The assumptions that can be made regarding the CALSC core subunit topology, stoichiometry, or preference for homo- or heterooligomers are naturally weakened. Structural characterization of other less-conserved CALS homologs (e.g. CALS10 or CALS12) would prove quintessential in answering these questions with more certainty. The CALS1 localization experiments relied on constitutive promoter-driven expression, which could possibly mask compartments with limited CALS1 presence or subtle fluctuations within PM domains. Further investigation of CALS1 subcellular localization under the native promoter will be the topic of future studies.

## Acknowledgment

This work was supported by Czech Science Foundation grant nr. 22-35680M (R.P.), nr. 26-23380N (P.C.) and nr. 23-04866S (M.W.). This study was also funded by project GA UK nr. 270823 (D.U.). Microscopy was performed at the Imaging Facility of the IEB CAS, supported by MEYS CR LM2023050 ‘Czech-BioImaging’ and IEB CAS. Computational resources used for structure modeling and molecular dynamics simulations were provided by the e-INFRA CZ project (ID: 90254), supported by the Ministry of Education, Youth and Sports of the Czech Republic. We thank the VIB proteomics core facility for performing the mass spectrometry data collection.

## Material and Methods

### Plant material and cultivation

*Arabidopsis thaliana* plants were grown in peat pellets and *in vitro*. For *in vitro* cultivation, seeds were surface-sterilized by chlorine gas, stratified for 2–3 days at 4°C and sown on vertical plates with agar medium (1 % [w/v] sucrose, 2.2 g/l MS salts [Sigma/Aldrich], 0.8 % [w/v] agarose, [pH 5.7]) in cultivation room under a photoperiod of 16 h light/8 h darkness and 21 °C. For propagation of plants, seedlings were transferred into peat pellets (Jiffy Products International) and grown as indicated above. For functional assays of PD permeability, surface-sterilized seeds were sown directly in Jiffy 7 peat pellets and plants were cultivated for 4 weeks under a short-day photoperiod (10 h/14 h light/dark regime) at 100-130 μE m^-2^ s^-1^, 22 °C and 70% relative humidity, and were watered with fertilizer-free distilled water as necessary. Genotypes used in this study were Col-0 (wild-type) and T-DNA insertional mutant lines *cals1* and *cals3* described previously (*cals1* - SAIL_1_H10^41^, *cals3* - SALK_068418^4^) and their double homozygote offspring line *cals1/cals3*.

Cells of tobacco line BY-2 (*Nicotiana tabacum* L. cv Bright-Yellow 2) were cultivated in darkness at 27 °C on an orbital incubator (150 rpm, orbital diameter 30 mm) in a liquid medium (3% [w/v] sucrose, 4.3 g/l Murashige and Skoog salts, 100 mg/l inositol, 1 mg/l thiamin, 0.2 mg/l 2,4-dichlorophenoxyacetic acid, and 200 mg/l KH_2_PO_4_ [pH 5.8]) and subcultured weekly. Stock BY-2 calli were maintained on media solidified with 0.6% (w/v) agar and subcultured monthly. BY-2 were transformed according to the previously published protocol^72^. Transgenic cells and calli were maintained on the same media supplemented with 50 μg/ml kanamycin or 50 μg/ml hygromycin. All chemicals were obtained from Sigma-Aldrich unless stated otherwise.

### Generation of constructs and transformation

To generate recombinant plasmids, genomic DNA of *Arabidopsis thaliana* (L.) Heynh *CALS1* (AT1G05570.1), *CALS1* coding sequence (CDS) and yeast strain *FKS1* (YLR342W) CDS were cloned into the Golden Gate^73^ and GoldenBraid^74^ compatible entry vectors.

The expression construct p35S::CALS1-GS^rhino^ was created using Golden Gate system from pGGK-AG^75^, pGG-A-p35S-B, pGG-B-Omega_leader-C, pGG-C-CALS1-D, pGG-D-GS^rhino^-E, pGG-E-t35S-F and pGG-F-tNOS-G. Similarly for the expression construct p35S::CALS1-TurboID, pGGK-AG, pGG-A-p35S-B, pGG-B-Linker-C, pGG-C-CALS1-D, pGG-D-2xG4S_TurboID_3xHA-E, pGG-E-tHSP18.2M-F, pGG-F-Linker-G were assembled. Next, both expression vectors were introduced into *Agrobacterium tumefaciens* strain C58C1 RifR (pMP90) and *Arabidopsis thaliana* PSB-D cell cultures were stably transformed according to the original protocol^76^.

The construct pUBQ10::mEGFP-CALS1 was assembled through Golden Gate system by recombining pGGK-AG, pGG-A-pUBQ10-B, pGG-B-mEGFP-C, pGG-C-CALS1-D, pGG-D-Linker-E, pGG-E-tHSP18-F and pGG-F-Linker-G. The expression construct was then introduced into the *Agrobacterium tumefaciens* strain GV2260 for BY-2 transformation. The same construct was also utilized to generate stable *Arabidopsis* lines through the floral dip method^77^.

The constructs pUBQ10::mEGFP-CALS3_Bag_ and pUBQ10::mScarletI-CALS3_Vta1_ were generated from the pGGK-AG, pGG-A-pUBQ10-B, pGG-B-mEGFP-C/pGG-B-mScarletI-C, pGG-C-CALS3_Vta1_-D/pGG-C-CALS3_Bag_-D, pGG-D-Linker-E, pGG-suE-tHSP18-F and pGG-F-Linker-G entry cassettes. These expression vectors were then introduced into the *A. tumefaciens* GV2260 strain and subsequently to *Arabidopsis thaliana* by floral dip.

For the yeast complementation assay, we adapted the respective CALS1 and FKS1 versions for the controlled homology recombination into the YPRCΔ15 locus of the yeast genome, according to the previously published protocol^74^. The expression cassette consisted of CALS1/FKS1 under the constitutive glyceraldehyde 3-phosphate dehydrogenase promoter (pTDH3) and terminator (TDH2t), tagged with 6xHA tag and mScarletI at the N and C termini, respectively.

### Affinity purification, TurboID, mass spectrometry and background filtering

For AP-MS and TurboID, *Arabidopsis* PSB-D cells were harvested three days after sub-culturing in fresh MSMO medium. For TurboID, 50 µM biotin was added to the medium 1 h prior biomass harvesting. The AP-MS experiments were performed in triplicate as described before^78^, with minor modifications for improved solubilization of CALS1 complexes through implementation of 1% (w/v) digitonin as detergent^79^. In short, a total protein extract was prepared in a digitonin-based extraction buffer (25 mM Tris-HCl pH 7.6, 15 mM MgCl_2_, 150 mM NaCl, 15 mM p-nitrophenyl phosphate, 60 mM β-glycerophosphate, 1% (w/v) digitonin, 0.1 mM Na_3_VO_4_, 1 mM NaF, 1 mM PMSF, 1 μM E64, EDTA-free Ultra Complete tablet (Roche), 5% (v/v) ethylene glycol and 0.1% benzonase). Protein complexes were trapped through the Protein G part of the GS^rhino^ tag by incubating 25 mg total protein extract per replicate for 45 min with 50 μL in-house prepared magnetic Immunoglobulin G (IgG) beads. After binding and removal of the non-bound fraction, beads were washed once with 500 μL extraction buffer, two times with 500 μL extraction buffer with 0.2% digitonin, once with 500 μL extraction buffer without digitonin and once with 800 μL 50 mM NH_4_HCO_3_ (pH 8.0). The wash buffer was removed, and beads were incubated in 50 μL 50 mM NH_4_HCO_3_ supplemented with 1 μg Trypsin/Lys-C (Promega) for 3 h at 37°C. Next, the digest was separated from the beads and incubated overnight with 0.5 μg Trypsin/Lys-C at 37°C. Finally, the digest was centrifuged at 20,800 rcf for 5 min, and supernatant was acidified to 1% (v/v) trifluoroacetic acid, desalted on C18 Omix tips (Agilent), dried in a SpeedVac and stored at −20 °C until MS analysis. TurboID-based proximity labelling experiments were carried out in triplicate following a previously published protocol^80^. In short, total protein extraction was performed using a denaturing extraction buffer (100 mM Tris-HCl pH 7.5, 2% (w/v) SDS, 8 M Urea) followed by removal of the free biotin with PD10 (Cytiva) columns. Biotinylated proteins were purified from the total protein extracts using Streptavidin Sepharose beads, followed by Trypsin/Lys-C digest and acid elution of biotinylated peptides.

For MS analysis, peptides were re-dissolved in 20 μl loading solvent A (0.1% TFA in water/ACN (98:2, v/v)) of which 2 μL was injected for LC-MS/MS analysis on an Ultimate 3000 RSLC nano LC (Thermo Fisher Scientific, Bremen, Germany) in-line connected to a Q Exactive mass spectrometer (Thermo Fisher Scientific). The peptides were first loaded on a PepMap^TM^ Neo Trap column (Thermo Fisher Scientific). After flushing from the trapping column, peptides were separated on a 50 cm μPAC™ column with C18-endcapped functionality (Pharmafluidics, Belgium) kept at a constant temperature of 50°C. Peptides were eluted by a non-linear gradient starting at 1% MS solvent B (0.1% FA in water/ACN, 20/80 (v/v)) and 99% MS solvent A (0.1% FA in water). For the gradient, MS solvent B reached 10% after 9 min, 33% at 28 min or at 32 min, 55% at 43 min, 70% at 48 min, followed by a 2 min wash with 70% MS solvent B and re-equilibration with 1% MS solvent B. Flow rate was set at 0.75 µL/min for 9 min and switched to 0.3 µL/min for the rest of the run. The mass spectrometer was operated in data-dependent mode, automatically switching between MS and MS/MS acquisition for the 5 most abundant peaks in each MS spectrum. Full-scan MS spectra (400–2,000 m/z) were acquired at a resolution of 70,000 in the Orbitrap analyzer after accumulation to a target value of 3,000,000 for a maximum of 80 ms. The 5 most intense ions above a threshold value of 13,000 were isolated with a width of 2 m/z for fragmentation at a normalized collision energy of 25% after filling the trap at a target value of 50,000 for maximum 80 ms. MS/MS spectra (200–2000 m/z) were acquired at a resolution of 17,500 in the Orbitrap analyzer. The polydimethylcyclosiloxane background ion at 445.12002 Da was used for internal calibration (lock mass).

For protein identification and filtering of non-specific background proteins, Thermo raw files were searched with Mascot (version 2.8.2, Matrix Science) vs the Araport11plus database as described before^81^. To determine specific proteins, a large dataset approach was followed^78,81^. For TurboID-MS, the large control PL-MS dataset consisted of 293 PL-MS experiments with bait proteins not known to be related to CALS1, and performed under similar conditions. For each identified protein a Normalized Spectral Abundance Factor (NSAF) was calculated. The Ln-transformed mean NSAF of all proteins identified in at least 2 replicates out of 3 were compared to the Ln-transformed mean NSAF of the same protein in the large control dataset by a two-tailed Student’s t-test. T-test p-values were -log_10_ transformed and infinite values were replaced by a constant -log_10_(p value) of 325. For each identified protein, an Affinity Enrichment Score (AES) = NSAF ratio (bait/control) x -log_10_(p-value) was calculated. Identifications with an AES >= 20 were retained. A similar approach was followed for AP-MS, but here two filters were utilized: one against the large GS^rhino^ dataset with baits proteins not known to be related to CALS1, and another against the GS^rhino^ dataset consisting of experiments with digitonin treatment only, to suppress digitonin-related background. The large GS^rhino^ control dataset consisted of 800 AP-MS and the digitonin GS^rhino^ dataset contained 74 AP-MS experiments. Identifications were considered significantly enriched if they passed one of the following criteria: (i) two-peptide identifications present in at least two out of three replicates are significantly enriched if they were found with less than three other bait-groups in the large control AP-MS dataset or if they were enriched with a mean NSAF ratio ≥10 AND a -log_10_(p-value) ≥ 10, or with a mean NSAF ratio ≥20 AND a -log_10_(p-value) ≥ 8, (ii) one-peptide identifications present in all three replicates, if they were found in the large control dataset with less than seven bait-groups and if they were significantly enriched with a mean NSAF ratio ≥10 AND a -Log10(p-value) ≥ 10.

The volcano plots of proteins from the MS dataset were generated using the software VolcanoseR (https://huygens.science.uva.nl/VolcaNoseR/).

### TAP purification combined with BS3 cross-linking and LC-MS/MS

For XL-MS, protein complexes were purified in triplicate by TAP, following a TAP cell culture protocol^82^ with minor adaptations, and purified proteins were on-bead crosslinked as described earlier^78,83^. Per replicate, proteins were extracted from 20 g CALS1-GS^rhino^ expressing cell suspension cells in AP-MS extraction buffer with 1% digitonin. During TAP, 300 mg total protein input was used per replicate and complexes were bound on 175 μl IgG and streptavidin beads during both binding steps. To remove non-specific background, IgG beads were washed two times with 1.75 mL extraction buffer with 1% digitonin and five times with 1.75 mL wash buffer (10mM Tris HCl pH 8.0, 150mM NaCl, 0.5mM EDTA, 1µM E64, 1mM PMSF, 5% (v/v) Ethylene glycol) supplemented with 0.2% digitonin, whereas streptavidin beads were washed sequentially with 5.25 mL wash buffer supplemented with 0.2% digitonin, 1.75mL wash buffer without detergent, and finally with 7 mL PBS. Next, streptavidin beads were resuspended in 175 μl PBS supplemented with 1.2mM BS3 (Thermo Fisher, cat. no. A39266) and crosslinked for 45 min at room temperature. After crosslinking, the reaction was quenched in 50 mM NH_4_HCO_3_ (pH 8.0) for 30 min at room temperature. Proteins were reduced in 5 mM dithiothreitol (DTT) and alkylated with 15 mM iodoacetamide. Next, beads were washed with 50 mM NH_4_HCO_3_ and incubated overnight at 37 °C with 1 μg Trypsin/Lys-C. The next day, an additional incubation was done with 0.5 μg Trypsin/Lyc-C. The digest was removed from the beads and desalted with Monospin C18 columns, dried using a SpeedVac and stored at −20 °C.

The peptides were re-dissolved in 20 µl loading solvent A (0.1% TFA in water/ACN (99.5:0.5, v/v)) of which 5 µl was injected for LC-MS/MS analysis on an Ultimate 3000 ProFlow nanoLC system in-line connected to a Q Exactive HF Biopharma mass spectrometer (Thermo). Trapping was performed at 20 μl/min for 4 min in loading solvent A on a 20 mm trapping column (made in-house, 100 μm internal diameter (I.D.), 5 μm beads, C18 Reprosil-HD, Dr. Maisch, Germany). The peptides were separated on an in-house produced column (75 µm x 250 mm), equipped with a laser pulled electrospray tip using a P-2000 Laser Based Micropipette Puller (Sutter Instruments), packed in-house with ReproSil-Pur basic 1.9 µm silica particles (Dr. Maisch). The column was kept at a constant temperature of 50°C. Peptides were eluted by a gradient reaching 26 % MS solvent B (0.1% FA in acetonitrile) after 135 min, 44% MS solvent B at 155 min, 56% MS solvent B at 160 min followed by a 5-minutes wash at 56% MS solvent B and re-equilibration with MS solvent A (0.1% FA in water). The flow rate was set to 250 nl/min. The mass spectrometer was operated in data-dependent mode, automatically switching between MS and MS/MS acquisition for the 16 most abundant ion peaks per MS spectrum. Full-scan MS spectra (375-1500 m/z) were acquired at a resolution of 60,000 in the Orbitrap analyzer after accumulation to a AGC target value of 3,000,000. The 16 most intense ions above a threshold value of 13,000 were isolated (isolation window of 1.5 m/z) for fragmentation at a normalized collision energy of 28% after filling the trap at a AGC target value of 100,000 for maximum 80 ms. MS/MS spectra (200-2,000 m/z) were acquired at a resolution of 15,000 in the Orbitrap analyzer. Precursor ions with unassigned, single and double charge states were excluded from fragmentation selection.

The raw files were processed with pFind3 (Chi et al., 2018) Open search against the Araport11plus database. The top 10 Arabidopsis proteins from the pFind search and Streptavidin were used to create a custom database that was used for searching crosslinked peptides in the pLink2 (version 2.3.11) program (Chen et al., 2019). Parameters were set as follows: flow type to conventional crosslinking with BS3, enzyme is Trypsin, with 3 missed cleavages allowed, peptide mass window is 600-6000, peptide length is 6-60, with precursor and fragment mass tolerances set to 20 ppm. Fixed modification was set to Carbamidomethylation of cysteine and variable modification was set to oxidation of methionine. Result filter tolerance was set to 10 ppm and FDR ≤ 5% at the PSM level.

### Gene ontology enrichment

Gene ontology (GO) enrichment analysis was performed using ShinyGO v0.8.2 (http://bioinformatics.sdstate.edu/go/). The significantly enriched hits from the MS analysis were searched against the Biological process gene set database with P-value cut-off of 0.05 and the top 30 pathways. The GO term network was visualized within Cytoscape v3.10.3^84^. The nodes were connected if they contained at least 20% identical GO terms. The node size indicated the gene count within the representative node, the node hue corresponded to the enrichment FDR of the GO term and the line thickness was relative to the number of shared genes between the two nodes.

### Subcellular localization analysis and StringDB annotation

For the subcellular localization annotation of the CALS1 interactome hits, we subjected the dataset to SUBA5 online tool^31^. Simultaneously we used StringDB^30^ (version 12.0) to create a functional protein association network. For the StringDB analysis, the default settings were used: the interaction score cutoff was set to 0.400 (medium) and evidence from all sources including experimental, co-expression and text mining was utilized. The networked proteins in the resulting dataset were clustered based on the SUBAcon consensus localization and visualized in Cytoscape.

### Split-ubiquitin assay, yeast complementation experiments, *in vivo* CALS activity assay

Putative interactors of CALS1 were validated by a split-ubiquitin system according to the previously published protocol^85^. Briefly, plasmids with the tested binding partners tagged with either C-or N-terminal moiety of ubiquitin (Cub and Nub, respectively) were cotransformed into THY.AP4 strain (genotype MATa ura3 leu2 lexA::lacZ::trp1 lexA::HIS3 lexA::ADE2) using the LiAc-PEG method^86^. As a control, prey plasmids containing NubG or wild type Nub were used as negative or positive controls, respectively. The selection was performed on a CSM medium lacking Leu, Trp and supplemented with 2% (w/v) glucose and 50 μM Met. After 2 days outgrowth, the colonies were resuspended in sterile milliQ water, diluted to OD_600_ of 1.0, 0.1 and 0.01 and plated on selective medium lacking Ade and His and supplemented with an increasing concentration of Met (50 - 750 μM). Colony growth at the respective dilutions was recorded after 3 days.

Plasmids containing the CALS1 or FKS1 modified by site-directed mutagenesis (SDM) at specific residues were linearized by *Not*I and introduced into the fks1p mutant strain BY21271 (genotype MATα aro7 can1 leu2 trp1 ura3 fks1::URA3; referred to as Δfks1)^65^, obtained from NBRP, Japan (JPNBRP202225). The transformation mixture was plated on YPD+200 µM G418 (Sigma-Aldrich). After two days, the transformed colonies were screened by colony PCR to confirm the DNA introduction.

To measure CALS enzymatic activity *in vivo*, we adapted the aniline blue fluorochrome-based method, originally shown in^87,88^. In detail, each Δfks1-complementing strain was grown overnight in YPD, after which it was pelleted at 1500 g and resuspended in milliQ water to OD_600_ 1.0. The cultures were then mixed with 4x diluted Aniline Blue Fluorochrome (Biosupplies Australia) in a ratio of 1:1, directly on the microscopic slide. The yeast cells were examined with an Zeiss LSM880 confocal microscope with C-Apochromat 40x/1.2 W objective; the fluorescence of the Aniline Blue was recorded with 405 nm Ex/410-585 nm Em settings at 4% laser intensity. The same imaging setup was used for all samples. To infer the β-1-3-glucan levels, the images were analyzed in Fiji software with the built-in Analyze particles tool at default settings (selection was restricted to live yeast cells of 15-60 µm^2^). The data were normalized to the mean Δfks1 background strain signal per each experiment; the differences between individual strains were analyzed by the Kruskal-Wallis test with Dunn post hoc test (p value < 0.05).

### Confocal microscopy

BY-2 suspension cells were imaged three days after subculturing. Cells were incubated in 4× diluted Aniline Blue Fluorochrome (Biosupplies Australia) for 5 min to stain callose, followed by co-staining with the plasma membrane (PM) marker FM4-64 (1 mM stock, Thermo Fisher Scientific) diluted 1:1000 for 5 min prior to observation. The stained BY-2 cells together with the BY-2 pUBQ10::mEGFP-CALS1 line were imaged using a Zeiss LSM 900 confocal microscope equipped with an Airyscan 2 detector and a C-Apochromat 40×/1.20 W objective in multiplex Airyscan mode to enable fast, high-resolution 3D acquisition laser intensity. Fine Z-stacks were acquired using the following settings: GFP (excitation 488 nm, emission 495–550 nm), FM4-64 and propidium iodide (PI) (excitation 561 nm, emission 555–620 nm), and Aniline Blue (excitation 405 nm, emission 410–585 nm). Raw data were processed using ZEN Black software (Carl Zeiss, Jena, Germany). Subsequent visualization and 3D analysis were performed in ZEN Blue.

Cotyledons and roots from 7 DAG stably transformed plants expressing mEGFP-CALS1 were imaged as described above. Root tip cells were stained with FM4-64 for 5 min prior to imaging.

For pit fields diameter analysis in cotyledon pavement cells, 7 DAG cotyledons were stained with PI (Biotium; 1 M stock in water). Pavement cells were imaged using the Zeiss LSM 900 with Airyscan 2 and the C-Apochromat 40×/1.20 W objective in multiplex Airyscan mode. Fine Z-stacks were acquired. Pit fields, visible as regions of reduced PI staining along anticlinal walls, were manually measured using ZEN Blue software with 3D and orthogonal view modes.

To visualize callose, 7 DAG roots were incubated for 10 min in 3× diluted Aniline Blue Fluorochrome. 14 DAG cotyledons were incubated for 1 h in undiluted Aniline Blue Fluorochrome and subsequently co-stained with PI for 25 min to reveal pit field regions. Samples were mounted directly on microscope slides and imaged using the Zeiss LSM 900 with Airyscan 2 in multiplex mode.

To discriminate the fluorescence origin of the aniline blue fluorochrome signal, the FM4-64 signal, and the mEGFP-CALS1 signal within PD pit fields, imaging was performed using a Mirava Polyscope system (Abberior) equipped with a 40×/1.15 W objective and time-gated detection.

Statistically significant differences between groups were determined using one-way ANOVA, with significance (p value < 0.05) indicated by different letters.

### Plasmodesmata permeability assay

For symplastic connectivity assessment, 4-5 weeks old *Arabidopsis thaliana* plants’ true leaves in similar developmental stage (lines Col-0, *cals1*, *cals3*, *cals1/3*) were bombarded with 1 nm gold particles (BioRad) coated with 5 µg of plasmid DNA (pB7WG2.0GFP – free GFP under 35S; pB7WG2.0RFPER – ER-targeted RFP under 35S, 35S::SP-RFP-KDEL^89^} resuspended in 100 µL of 100% ethanol, using the biolistic PDS-1000/He particle delivery system (BioRad) with rupture discs for 1100 psi. 1 h after bombardment leaves were infiltrated with 1 mM salicylic acid or dH_2_O (mock). Leaves were kept in Petri plates with 1/2MS medium containing 0.7% (w/v) agar at room temperature overnight before imaging.

Images were taken ∼24 h post-bombardment, each bombardment site considered as an independent value, 5-10 bombardment sites per leaf counted. For imaging, RFP was excited with a 561-nm DPSS laser and emission collected at 600 to 640 nm, while GFP was excited with a 488-nm argon laser and emission collected at 505 to 530 nm using a Plan-Apochromat 10x/0.45 M27 objective on Zeiss LSM880 confocal imaging system.

Data from 4 repeats were analysed together (n=84-153 in total). One repeat consisted of 4 plants per genotype, 3 leaves per plant were sampled, bombarded, treated with SA and analyzed (at least 5 bombardment sites per leaf). Within each repeat, values were normalized to the average movement for mock in the experiment and put together as a “Relative movement”. Letter annotation was determined from one-way ANOVA with Tukey’s multiple comparisons test (p value < 0.05) results, and confirmed by bootstrappin^909^.

### Protein isolation, cell fractionation

BY-2 plasma membrane fractions were prepared from 105 ml of 2 day old culture expressing mEGFP-CALS. Excess growth medium was filtered and the residual cells were sonicated by 3x60 s bursts in homogenization buffer (50 mM HEPES, 400 mM sucrose, 100 mM KCl, 100 mM MgCl_2_, pH 7.5) containing cOmplete Protease Inhibitors cocktail (Roche). The homogenate was filtered through a mesh cloth and sedimented at 6000 g (10 min, 4°C). The supernatant containing the extracted proteins was transferred to a 20 ml ultracentrifuge cuvettes and subsequently centrifuged at 150 000 g (60 min, 4°C) to separate the cytosolic (supernatant) and microsome-containing (pellet) fractions. The obtained pellet was resuspended in 2.5 ml phosphate buffer (5 mM KH_2_PO_4_, 5 mM Na_2_HPO_4_, pH 7.8) and homogenized in a Potter-Elvehjem homogenizer. 2 g of the microsomal fraction was loaded on top of a glass cuvette containing two-phase PEG-dextran partitioning gradient. The phases were mixed by swing movement and left at 4°C overnight to allow the phase separation. The PM-containing top fraction was transferred to a second cuvette containing an identical gradient, mixed thoroughly and sedimented at 520 g (5 min, 4°C). The top fraction was transferred to an ultracentrifugation cuvette and centrifuged at 150 000 g (60 min, 4°C). Finally, the supernatant was removed and the purified PM fraction was resuspended in 5 mM of phosphate buffer.

A similar approach was adopted for *Arabidopsis thaliana* stable lines expressing mEGFP-CALS_FL_, -CALS_Bag_ or mScarletI-CALS_Vta1_. Briefly, approximately 0.5 g of 2 week old seedlings grown on 1/2 MS plates on top of a nylon mesh were ground on ice with a pre-chilled homogenization buffer and subjected to the steps mentioned above.

To extract yeast membrane fractions, 250 ml of turbid YPD-grown yeast culture was sedimented at 1500 g. The pelleted culture was washed in milliQ water and spun down once again. The resulting pellet was resuspened in 2.5 ml of homogenization buffer, transferred in tiny drops into a mortar filled with liquid nitrogen and ground to a fine powder. The same approach as demonstrated for plant samples was followed in the subsequent steps.

### SDS-PAGE, Blue native PAGE, Western blotting and immunodetection

Protein samples isolated from the respective plant or yeast tissues were analyzed by Western blotting (WB) and immunodetection. In detail, the samples were separated on 4-20% stain-free polyacrylamide SDS-gels (Bio-Rad) at 180 V and transferred to a Trans-Blot Turbo PVDF membrane (Bio-Rad) at 12 V. Membranes were rinsed in 1xTBS-T buffer, blocked in 5% non-fat milk overnight and incubated with primary antibodies for 60 min. Subsequently, the excess antibody was removed by washing the membrane thrice in TBS-T and incubated with a HRP-conjugated secondary antibody (α-mouse; Promega) for 60 min. All antibodies were diluted in 5% milk/TBS-T. The chemiluminescent signal was detected following Radiance Plus (Azure) coincubation by ChemiDoc Imaging System (Bio-Rad).

For the investigation of non-denatured protein complexes, blue native PAGE was employed according to the original protoco^910^. For the first dimension, previously obtained plasma membrane fractions (40 µl, 10 µg/µl) in resuspension buffer (10% glycerol, 0.05 M NaCl, 0.05 M imidazole, pH 7.0) were buffer-exchanged by centrifugation at 27,460 × g (15 min, 4 °C). Then, pellets were resuspended in solubilization buffer B (50 mM imidazole, 500 mM 6-aminohexanoic acid, 1 mM EDTA, 0.5% n-dodecyl-β-D-maltoside [DDM]), incubated for 15 min at 4 °C, and centrifuged again at 27,460 g (30 min, 4 °C). Supernatants were supplemented with 0.06 % (w/v) Coomassie Blue G-250 (Bio-Rad) and 0.125 % (v/v) glycerol prior to Blue Native PAGE on 4–15% Stain-Free gels (Bio-Rad) at (180 V, 4 °C). Proteins separated in the first dimension were either transferred to a membrane for WB or used for the second dimension. In the latter case, selected gel strips were denatured (2% SDS, 50 mM Tris-HCl, pH 6.8), placed horizontally on self-cast gradient gels (8–4%), separated by SDS-PAGE and analyzed by WB.

### CALS *in vitro* enzymatic assay

The enzymatic activity of the isolated CALS-containing fraction was tested as described previousl^921^. In brief, 2.5 µg of PM fraction was added to the microtiter plate well containing 50 µl of the reaction mixture (50 mM HEPES, 0.01% (w/v) digitonin, 1 mM CaCl_2_, 10 mM cellobiose, 0.4 mM UDP-glucose, pH 7.3) and incubated at 25°C for 30 min. The reaction was stopped by the addition of 10 µl 6 N NaOH and the reaction product was solubilized by heating the plate to 80°C. Afterwards, the released glucan was incubated with 210 µl of 0.1 % (w/v) aniline blue at 50°C for 30 min. The end-point product was detected in a spectrophotometer (λ_ex_=400 nm, λ_em_=460 nm).

### AlphaFold model prediction and scoring

Multiple neural network-based model prediction softwares were used in this study. For all predictions of proteins in a monomer state with no ligands, ColabFold2 modification of the AlphaFold2 (AF2) was employe^932^. To determine the binding mode between a monomeric protein and ligands, we used AlphaFold3 (AF3)^943^. Structures of oligomeric assemblies were predicted using AF2-MM3^53^. The quality and local confidence of the obtained models were assayed based on the pLDDT score, while the global confidence and interaction between subunits was assessed by the ipTM and predicted alignment error (PAE) metrics.

In all cases, the full length version of the CALS1 protein was utilized except for the 5x and 6x assemblies of CALSC, where Vta1 and Bag domains were omitted to fit within the computational limits. We noted that the presence of Vta1 and Bag domains was not necessary for the correct inference of the CALSC overall topology

### Molecular dynamics simulations and trajectory analysis

All molecular dynamics (MD) simulations were executed using the GROMACS software version 2020.^954^. For the simulation of CALS1, the atomic representation AF2 structures were converted to the CHARMM36m forcefield in the CHARMM-GUI v3.8 web serve^965^. Detailed simulation parameters for all runs are in **Supplementary Data**.

### Functional domain classification, structural similarity search, transport channel analysis

The previously published dataset of the CALS homologs^40^ was subjected to the InterPro web tool for the annotation of functional domain^976^.

For the analysis of CALS amino acids evolutionary conservation, the predicted structure was used as an input for the ConSurf web serve^987^. A custom MSA obtained from^40^, was utilized to direct the search across CALS homologs from plant and yeast.

Using the structural delineation of the CALS domains, the structure of the Bag domain was used for DALI^54^ and Foldseek^55^ structural similarity searches.

To predict the putative transport channels inside CALS, we utilized the Caver web 2.0 tool^69^. As an input, we used the AF2-predicted model fitted with the published ꞵ -1,3-glucan experimental structure^71^. Two timepoints at the start and end of a 500 ns MD simulation (see below) were used to determine the closed and open states of CALS. For all predictions, we used the default settings, with the exception of the tunnel starting point, which was set at the CALS active site center. Out of the calculated tunnels, only those which passed through the TM helix region were considered. For further analyses, the channel with the highest throughput metric was chosen.

---

**Fig. S1:**
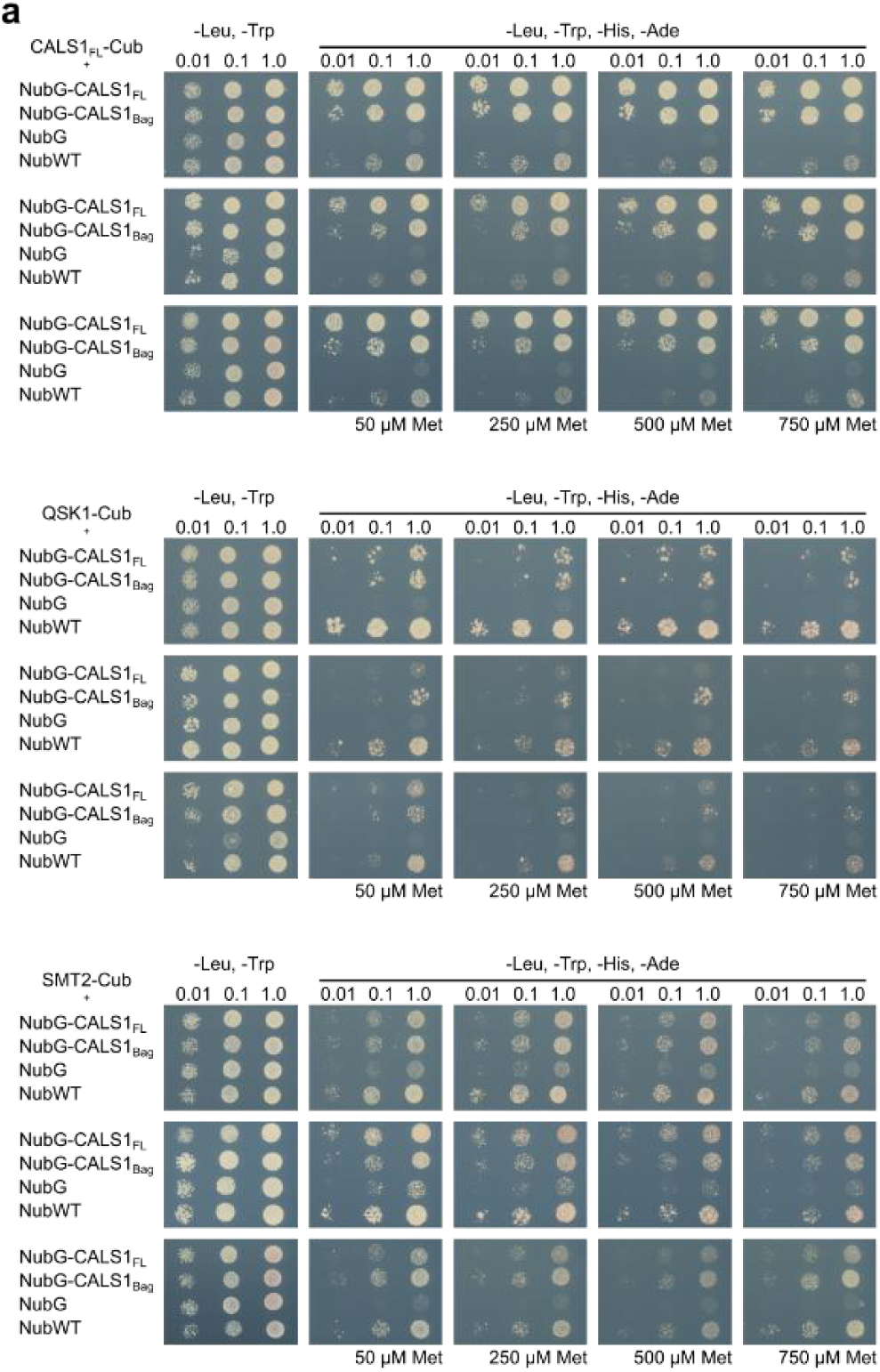
Validation of selected CALS1 interactors by split ubiquitin. (**a**) The interaction of bait and prey proteins tagged with Cub and Nub, respectively, was tested using the dilution spot assay at various auxotrophic conditions. The growth of the colonies correlates with the strength of the interaction.

**Fig. S2:**
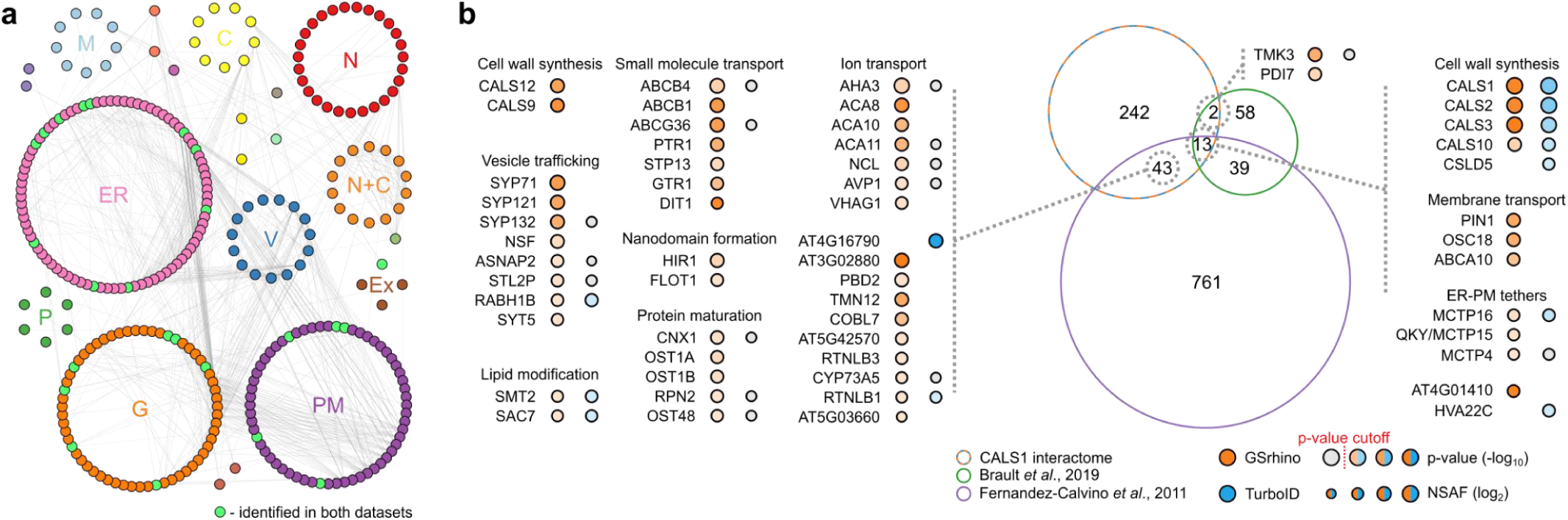
CALS1 interactome analyses. (**a**) Proteins within the AP-MS and TurboID datasets were clustered based on their SUBAcon consensus localization. Each node represents one protein, the light green indicates hits identified by both approaches. The nodes are connected according to the StringDB database of protein-protein interactions; each connection corresponds to one binary interaction. (**b**) Comparison of the CALS1 dataset with previously published plasmodesmatal proteomes. The proteins within each overlapping region were grouped based on their proposed biological function. The NSAF values obtained by the AP-MS and/or TurboID experiments are provided.

**Fig. S3:**
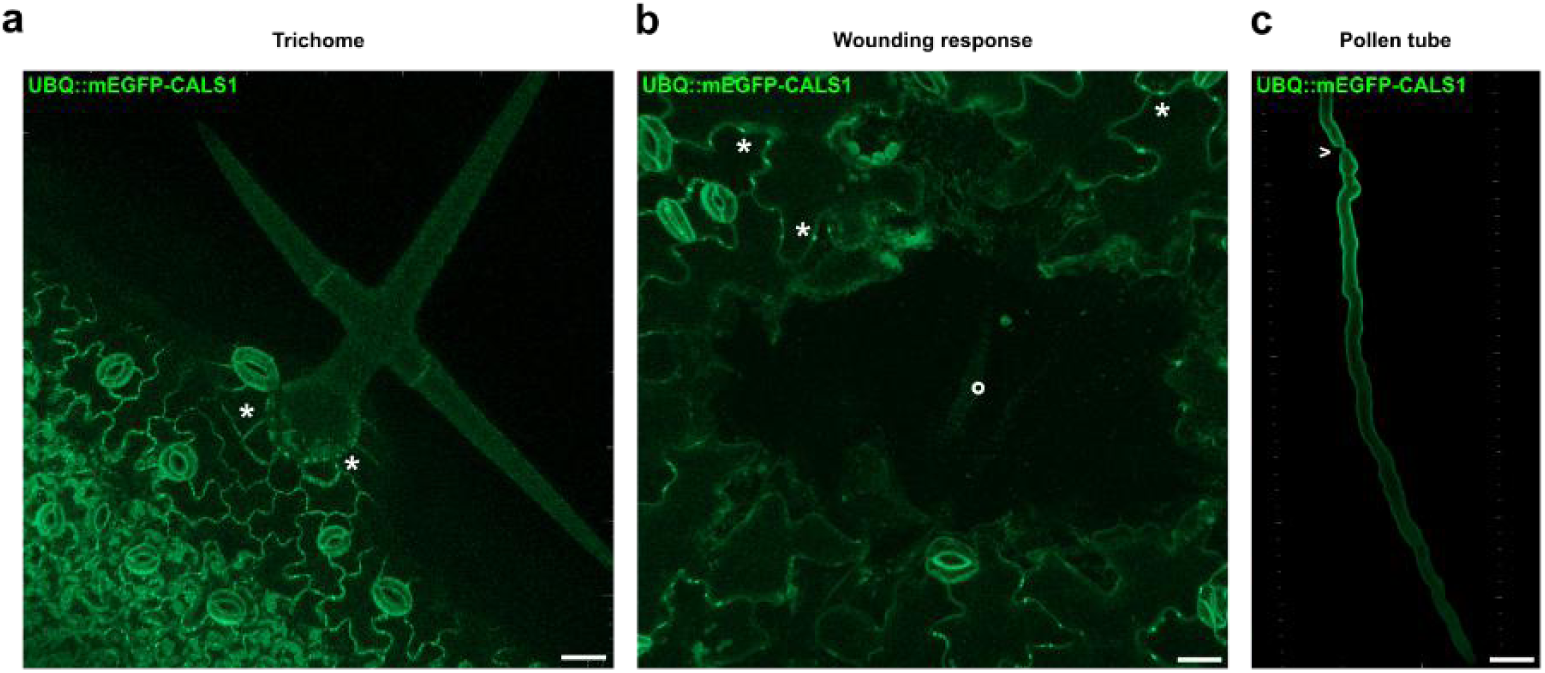
CALS1 subcellular localization *in planta*. *Arabidopsis thaliana* stable lines expressing mEGFP-CALS1. (**a**) In first leaf trichomes, signal is restricted to plasmodesmata (PD) (*) at the trichome base, with no apparent signal in place of the Ortmannian ring. (**b**) After wounding, CALS1 remains at the PD and does not relocalize to the wounding site (°) periphery (14 DAG cotyledon). (**c**) In the pollen tube, CALS is predominantly localized to the shank plasma membrane, mEGFP-CALS1 signal is absent in the pollen tube tip. No signal enrichment is observable in the callose plug region (>). Scale bar = 20 μm.

**Fig. S4:**
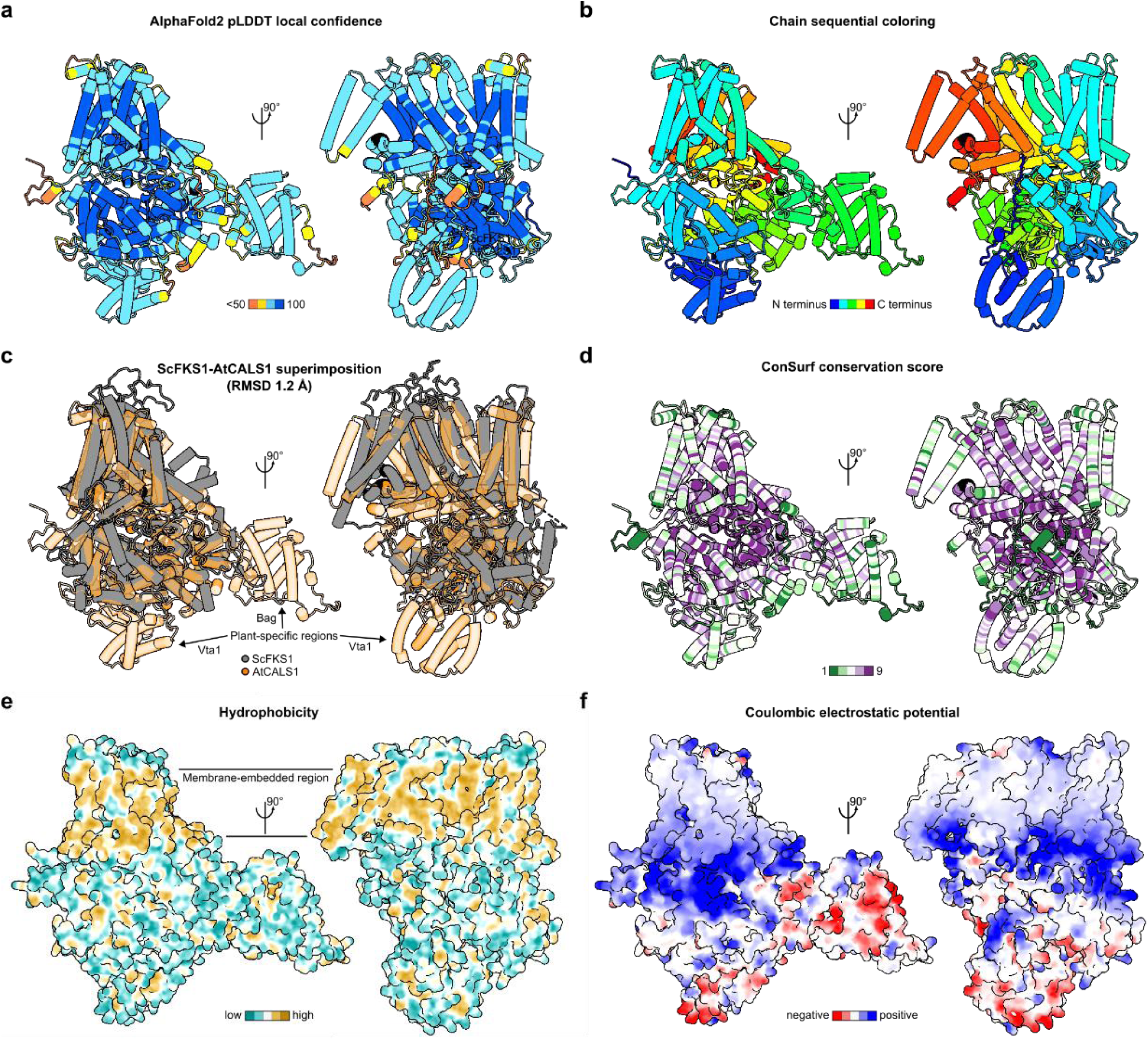
Overview of the CALS1 structure parameters. (**a**) Local model confidence, shown by pLDDT score. The most confident domains (dark blue) are present in the central core and the transmembrane (TM) helices. CALS1 contains only a small number of poorly scoring regions at the periphery of the structure. (**b**) CALS1 chain colored sequentially by a rainbow palette. (**c**) Superimposition of CALS1 with the budding yeast ScFKS1 structure (PDB ID 7YUY). The two related proteins are structurally to a large extent similar. The added regions present in CALS1 correspond to the Vta1 and Bag domains. (**d**) Evolutionary conservation of the individual residues, represented by the Consurf score. The central core is highly conserved, compared to the outer parts of the protein. The least conserved regions are the loops at the N and the C termini. (**e**) Surface representation of the CALS1, colored according to the molecular lipophilicity potential. Lipophilic residues (light brown) are concentrated within a single zone, forming the integral membrane part. The hydrophilic residues (cyan) encompass the section in contact with the cytoplasm. (**f**) Surface representation of the CALS1, colored by the Coulombic electrostatic potential. Several positively charged areas (blue) are present within the structure.

**Fig. S5:**
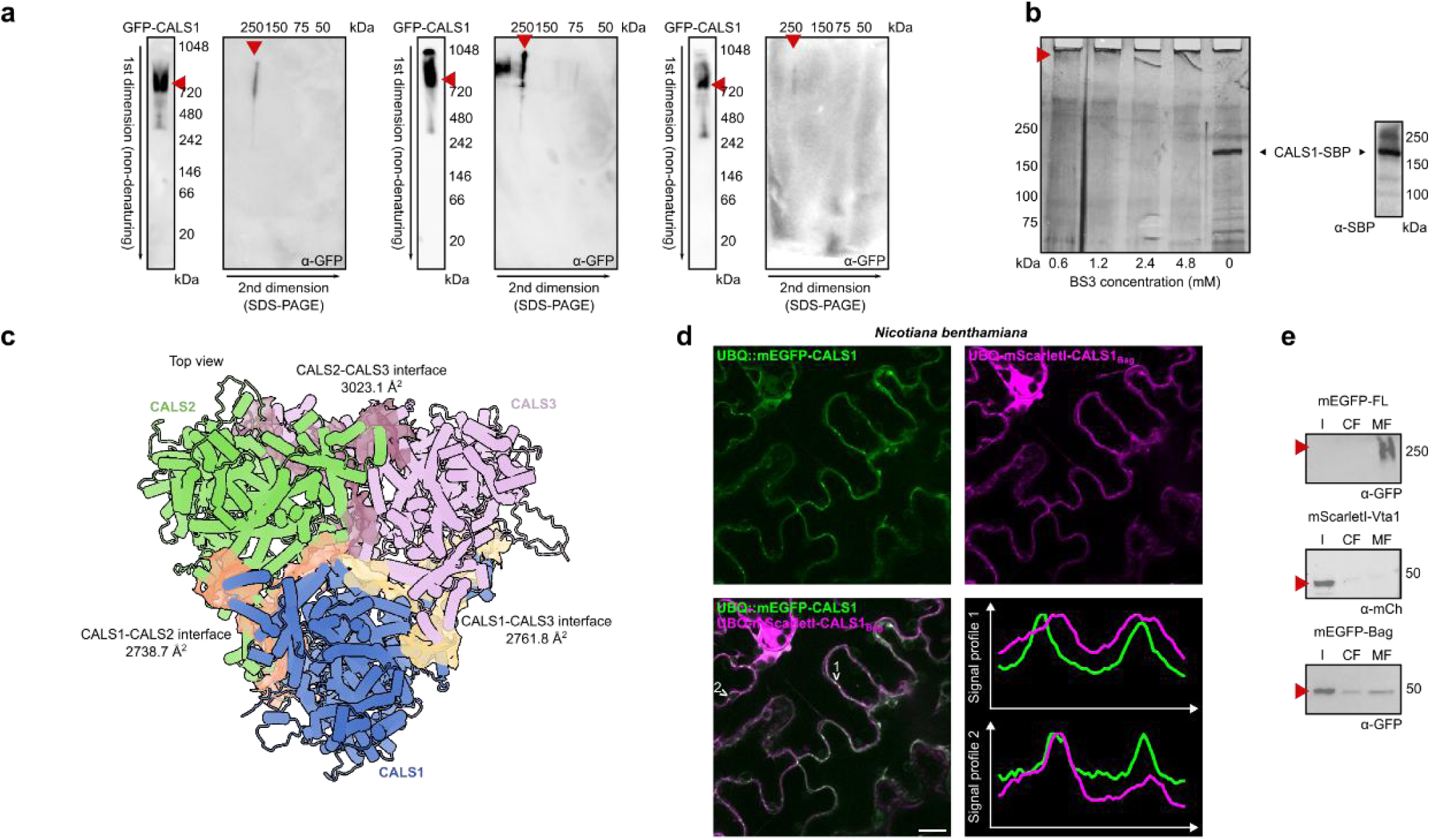
CALS oligomerization data. (**a**) Blue native PAGE of mEGFP-AtCALS1 expressed in tobacco BY-2 cell culture. The red triangle highlights bands corresponding to the trimeric (1st dimension) or denatured (2nd dimension) form of protein. (**b**) SDS-PAGE of CALS1-SBP expressed in Arabidopsis cell culture. Samples were treated on-beads with a BS3 crosslinker of increasing concentrations (0-4.8 mM). The red triangle shows the crosslinked protein, unable to penetrate the gel. Presence of protein in the sample was validated by WB. (**c**) Overview of the CALS heterotrimer prediction by AF2-MM3. The buried solvent-accessible surface area is highlighted with burgundy, orange and beige colors. (**d**) Colocalization analysis of CALS Bag domain. mEGFP-CALS1FL and mScarletI-CALS1Bag were transiently expressed in *N. benthamiana* leaves. The punctate pattern at the plasma membrane can be observed for both proteins. Scale bar = 20 μm. (**e**) Western blotting of samples from the cell fractionation of 14 days after germination Arabidopsis thaliana seedlings expressing different CALS domains. The full length (FL) protein is enriched in the microsomal fraction (MF); Bag domain exhibits populations both in cytosolic fraction (CF) and MF; Vta1 remains predominantly in CF with a small portion present in the MF.

**Fig. S6:**
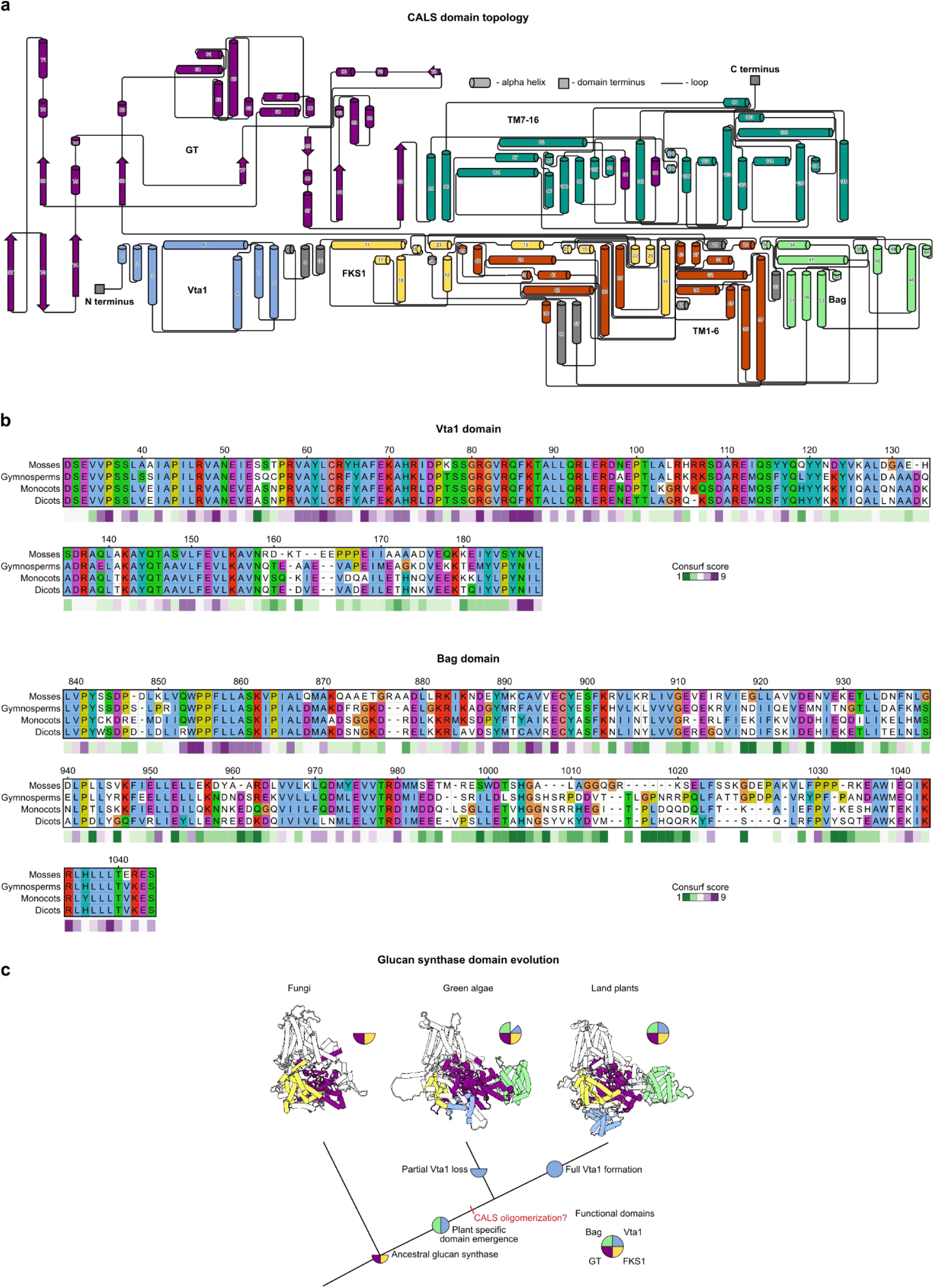
CALS functional domains and their evolutionary context. (**a**) 2D diagram of the CALS functional domains topology. Vta1 and Bag form independent regions, while FKS1 and Glycosyl transferase (GT) are interspersed within the complex structure. The mutual contact between the TM1-6 and TM7-16 is demonstrated. (**b**) Multiple sequence alignment (MSA) of Vta1 and Bag domains from selected plant species. The Consurf score highlighting the evolutionary conservation of the residues is provided below the MSA. (**c**) Suggested model of the glucan synthase domain evolution. The common ancestor of plants and fungi contained only the FKS1 and GT domains, while the plant specific Vta1 and Bag domains were acquired after the split of fungi and plants, possibly underpinning their role in CALS oligomerization. While the Bag domain is conserved within the plant kingdom, Vta1 underwent secondary reduction/loss in green algae.

**Fig. S7:**
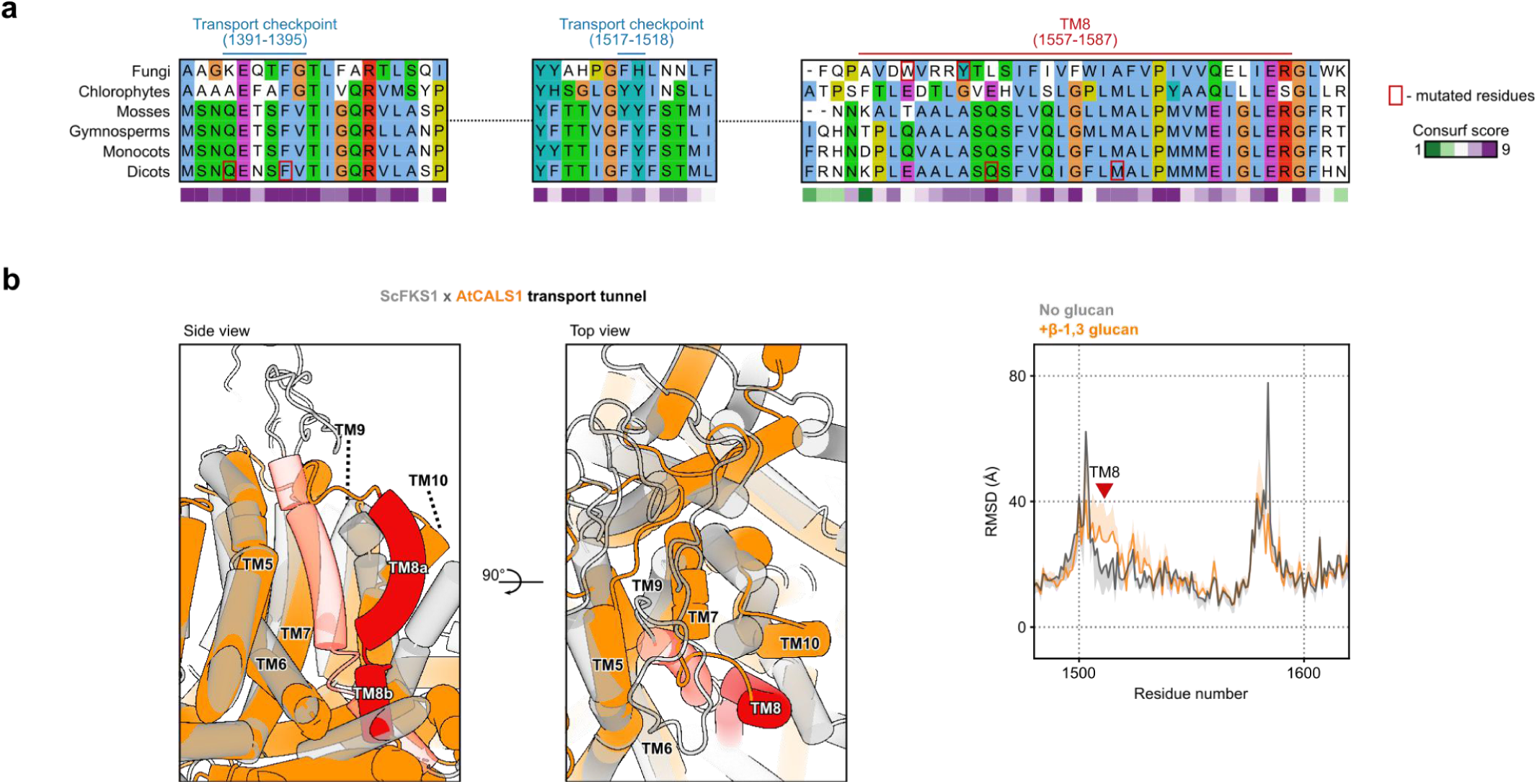
Evolutionary context of CALS translocation tunnel. (**a**) Multiple sequence alignment (MSA) of key transport regions from CALS homologs of selected species. The evolutionary conservation of the tunnel lining residues is represented by the ConSurf score below the MSA. Residues selected for mutation analysis are highlighted by a red square. (**b**) Superimposition of yeast (grey) and plant (orange) transporting regions. The TM8 helix is annotated in red. (**c**) Per residue RMSD between CALS structures containing glucan or no glucan after 500 ns molecular dynamics simulation. The position of TM8 is shown with a red triangle.

**Fig. S8:**
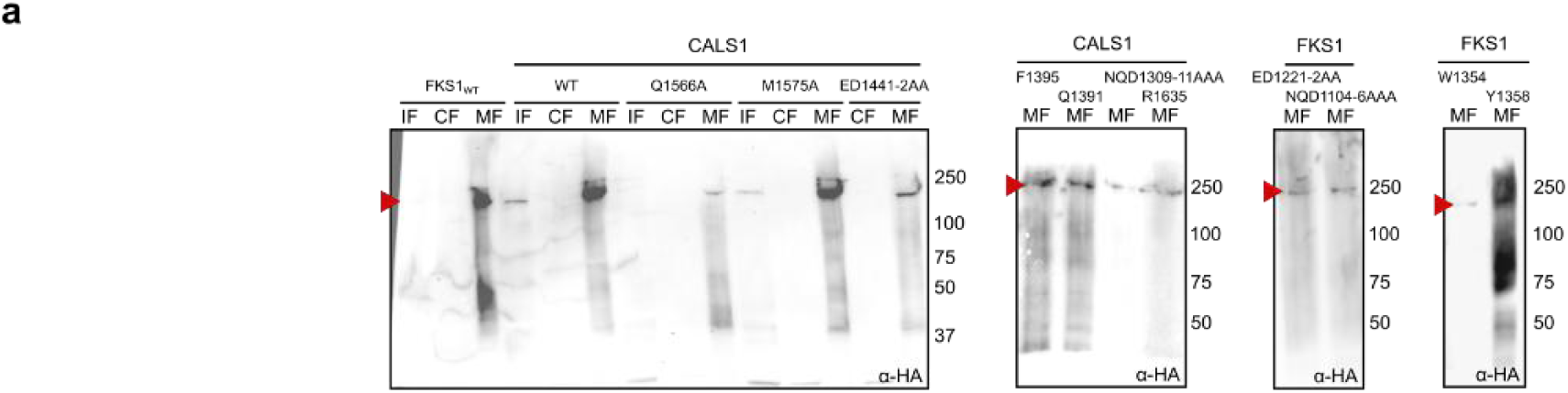
Validation of yeast complementation lines by Western blot. (**a**) Subcellular fractions separated by SDS-PAGE. The enriched bands (arrows) correspond to the size of CALS1 or FKS1. Input fraction (IF), cytosolic fraction (CF), microsomal fraction (MF).

